# Linked dimensions of psychopathology and connectivity in functional brain networks

**DOI:** 10.1101/199406

**Authors:** Cedric Huchuan Xia, Zongming Ma, Rastko Ciric, Shi Gu, Richard F. Betzel, Antonia N. Kaczkurkin, Monica E. Calkins, Philip A. Cook, Angel Garcia de la Garza, Simon Vandekar, Tyler M. Moore, David R. Roalf, Kosha Ruparel, Daniel H. Wolf, Christos Davatzikos, Ruben C. Gur, Raquel E. Gur, Russell T. Shinohara, Danielle S. Bassett, Theodore D. Satterthwaite

## Abstract

Neurobiological abnormalities associated with psychiatric disorders do not map well to existing diagnostic categories. High co-morbidity and overlapping symptom domains suggest dimensional circuit-level abnormalities that cut across clinical diagnoses. Here we sought to identify brain-based dimensions of psychopathology using multivariate sparse canonical correlation analysis (sCCA) in a sample of 663 youths imaged as part of the Philadelphia Neurodevelopmental Cohort. This analysis revealed highly correlated patterns of functional connectivity and psychiatric symptoms. We found that four dimensions of psychopathology — mood, psychosis, fear, and externalizing behavior — were highly associated (*r*=0.68-0.71) with distinct patterns of functional dysconnectivity. Loss of network segregation between the default mode network and executive networks (e.g. fronto-parietal and salience) emerged as a common feature across all dimensions. Connectivity patterns linked to mood and psychosis became more prominent with development, and significant sex differences were present for connectivity patterns related to mood and fear. Critically, findings replicated in an independent dataset (n=336). These results delineate connectivity-guided dimensions of psychopathology that cut across traditional diagnostic categories, which could serve as a foundation for developing network-based biomarkers in psychiatry.

## INTRODUCTION

Psychiatry relies on signs and symptoms for clinical decision making, while other branches of medicine are transitioning to the use of biomarkers to aid in diagnosis, prognosis, and treatment selection. The search for biomarkers in psychiatry has intensified,^1^ and it is increasingly recognized that existing clinical diagnostic categories could hinder this effort, as they do not pair well with distinct neurobiological abnormalities.^2–4^

The high co-morbidity among psychiatric disorders exacerbates this problem.^5^ Furthermore, studies have demonstrated common structural, functional, and genetic abnormalities across psychiatric syndromes, potentially explaining such co-morbidity.^6–10^ This body of evidence underscores the lack of direct mapping between clinical diagnostic categories and the underlying pathophysiology, potentially leading to dramatic changes to treatment strategies for psychiatric disorders.

This context has motivated the development of the National Institute of Mental Health’s Research Domain Criteria, which seek to construct a biologically-grounded framework for neuropsychiatric diseases.^11,12^ In such a model, the symptoms of individual patients are conceptualized as the result of mixed dimensional abnormalities of specific brain circuits. While such a model system is theoretically attractive, it has been challenging to implement in practice due to both the multiplicity of clinical symptoms and the many brain systems implicated in psychiatric disorders.^13,14^

Network neuroscience is a powerful approach for examining brain systems implicated in psychopathology.^15–17^ One network property commonly evaluated is its community structure, or modular architecture. A network module (also called a sub-network or a community) is a group of densely interconnected nodes, which may form the basis for specialized sub-units of information processing.

Converging results across data sets, methods, and laboratories provide substantial agreement on large-scale functional brain modules such as the somatomotor, visual, default mode, and fronto-parietal control networks.^18–26^ Furthermore, multiple studies documented abnormalities within this modular topology in psychiatric disorders.^16,27, 28^ Specifically, evidence suggests that many psychiatric disorders are associated with abnormalities in network modules subserving higher-order cognitive processes, including the default mode and fronto-parietal control networks.^29^

In addition to such module-specific deficits, studies in mood disorders,^30–32^ psychosis,^28,33–35^ and other disorders^36,37^ have reported abnormal interactions *between* modules that are typically segregated from each other at rest. This is of particular interest as modular segregation of both functional^19,38, 39^ and structural^40^ brain networks is refined during adolescence, a critical period when many neuropsychiatric disorders emerge. Such findings have led many disorders to be considered “neurodevelopmental connectopathies.”, ^41–44^ Describing the developmental substrates of neuropsychiatric disorders is a necessary step towards early identification of at-risk youth, and might ultimately allow for interventions that “bend the curve” of maturation to achieve improved functional outcomes.^45^

Despite the increasing interest in describing how abnormalities of brain network development lead to the emergence of neuropsychiatric disorders, existing studies have been limited in several respects. First, most adopted either a categorical case-control approach, or only examined a single dimension of psychopathology. Second, especially in contrast to adult studies, existing work in youth has often used relatively small samples (e.g. dozens of participants). While multivariate techniques could allow examination of both multiple brain systems and clinical dimensions simultaneously, such techniques usually require large samples.

In the current study, we sought to delineate functional network abnormalities associated with a broad array of psychopathology in youth. We capitalized on a large sample of youth from the Philadelphia Neurodevelopmental Cohort (PNC)^46^ applying a recently-developed machine learning technique called sparse canonical correlation analysis (sCCA).^47^ As a multivariate method, sCCA is capable of discovering complex linear relationships between two high-dimensional datasets.^48,49^ Here, we used sCCA to delineate linked dimensions of psychopathology and functional connectivity. As described below, we uncovered dimensions of dysconnectivity that were highly correlated with specific, interpretable dimensions of psychopathology. We found that each psychopathological dimension was associated with a pattern of abnormal connectivity, and that all dimensions were characterized by decreased segregation of default mode and executive networks (fronto-parietal and salience). These network features linked to each dimension of psychopathology showed expected developmental changes and sex differences. Finally, our results were replicated in an independent dataset.

## RESULTS

We sought to delineate multivariate relationships between functional connectivity and psychiatric symptoms in a large sample of youth. To do this, we used sCCA, an unsupervised learning technique that seeks to find correlations between two high-dimensional datasets.^47^ In total, we studied 999 participants ages 8-22 who completed both functional neuroimaging and a comprehensive evaluation of psychiatric symptoms as part of the PNC.^46,50^ We divided this sample into discovery (n=663) and replication datasets (n=336) that were matched on age, sex, race, and overall psychopathology (**Supplementary Fig. 1** and **Supplementary Table 1**). Following pre-processing using a validated pipeline that minimizes the impact of in-scanner motion,^51^ we constructed subject-level functional networks using a 264-node parcellation system^19^ that includes an *a priori* assignment of nodes to network communities (**Fig. 1a-c**. e.g. modules or sub-networks; see Online Methods). Prior to analysis with sCCA, we regressed age, sex, race, and motion out of both the connectivity and clinical data to ensure that these potential confounders did not drive results. As features that do not vary across subjects cannot be predictive of individual differences, we limited our analysis of connectivity data to the top 10 percent most variable connections (ranked by median absolute deviation, see Online Methods and **Supplementary Fig. 2)**. The input data thus consisted of 3410 unique functional connections (**Fig. 1b**) and 111 clinical items (**Fig. 1c** and **Supplementary Table 3**). Using elastic net regularization (*L*1 + *L*2), sCCA was able to obtain a sparse and interpretable model while minimizing over-fitting (**Fig. 1d** and **Supplementary Fig. 3**; see Online Methods). Ultimately, sCCA identified specific patterns (“cnonical variates”) of functional connectivity that were linked to distinct combinations of psychiatric symptoms.

**Figure 1.**
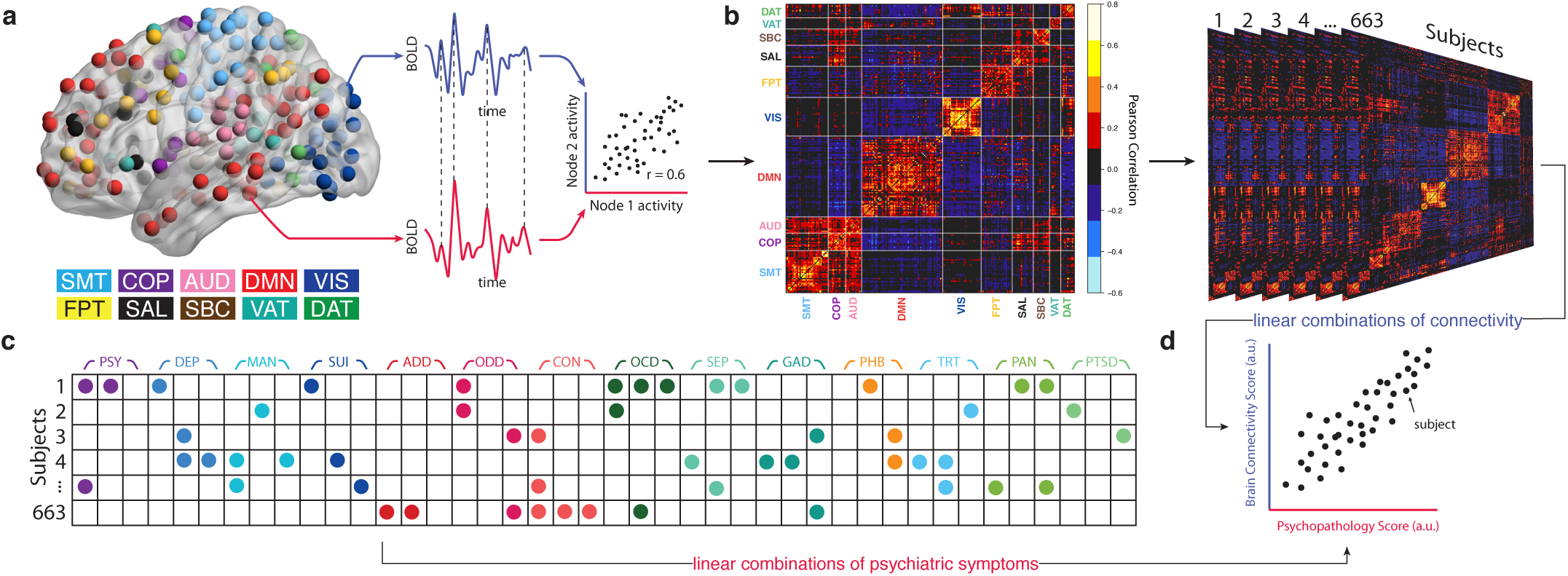
Schematic of sparse canonical correlation analysis (sCCA). **(a)** Resting-state fMRI data analysis schematic and workflow. After preprocessing, blood-oxygen-level dependent (BOLD) signal time series were extracted from 264 spherical regions of interest distributed across the cortex and subcortical structures. Nodes of the same color belong to the same *a priori* community as defined by Power et al.^19^ **(b)** Awhole-brain, 264 *×* 264 functional connectivity matrix was constructed for each subject in the discovery sample (n =663 subjects). **(c)** Item-level data from a psychiatric screening interview (111 items, based on K-SADS^81^) were entered into sCCA as clinical features. **(d)** sCCA seeks linear combinations of connectivity and clinical symptoms that maximize their correlation. *A priori* community assignment: SMT: somatosensory/motor network; COP: cingulo-opercular network; AUD: auditory network; DMN: default mode network; VIS: visual network; FPT: fronto-parietal network; SAL: salience network; SBC: subcortical network; VAT: ventral attention network; DAT: dorsal attention network; Cerebellar and unsorted nodes not visualized. Psychopathology domains: PSY: psychotic and subthreshold symptoms; DEP: depression; MAN: mania; SUI: suicidality; ADD: attention-deficit hyperactivity disorder; ODD: oppositional defiant disorder; CON: conduct disorder; OCD: obsessive-compulsive disorder; SEP: separation anxiety; GAD: generalized anxiety disorder; PHB: specific phobias; TRT: mental health treatment; PAN: panic disorder; PTSD: post-traumatic stress disorder.

### Multivariate analysis reveals linked dimensions of psychopathology and connectivity

Based on the scree plot of covariance explained (**Fig. 2a**), we selected the first seven canonical variates for further analysis. Significance of each of these linked dimensions of symptoms and connectivity was assessed using a permutation test (see Online Methods and **Supplementary Fig. 4)**; False Discovery Rate (FDR) was used to control for type I error rate due to multiple testing. Of these seven cannonical variates, three were significant (Pearson correlation *r* = 0.71, *P*_*FDR*_ < 0.001; *r* = 0.70, *P*_*FDR*_ < 0.001, *r* = 0.68, *P*_*FDR*_ < 0.01, respectively) (**Fig. 2b**), with the fourth showing a trend towards significance (*r* = 0.68, *P*_*FDR*_ = 0.07,*P*_*uncorrected*_ = 0.04). Notably, these results were robust to many different methodological choices, including the number of features entered into the initial analysis (**Supplementary Fig. 5a**), the parcellation system (**Supplementary Fig. 5b**), and the use of regularization with elastic net versus data reduction with principal component analysis (**Supplementary Fig. 5c**).

**Figure 2.**
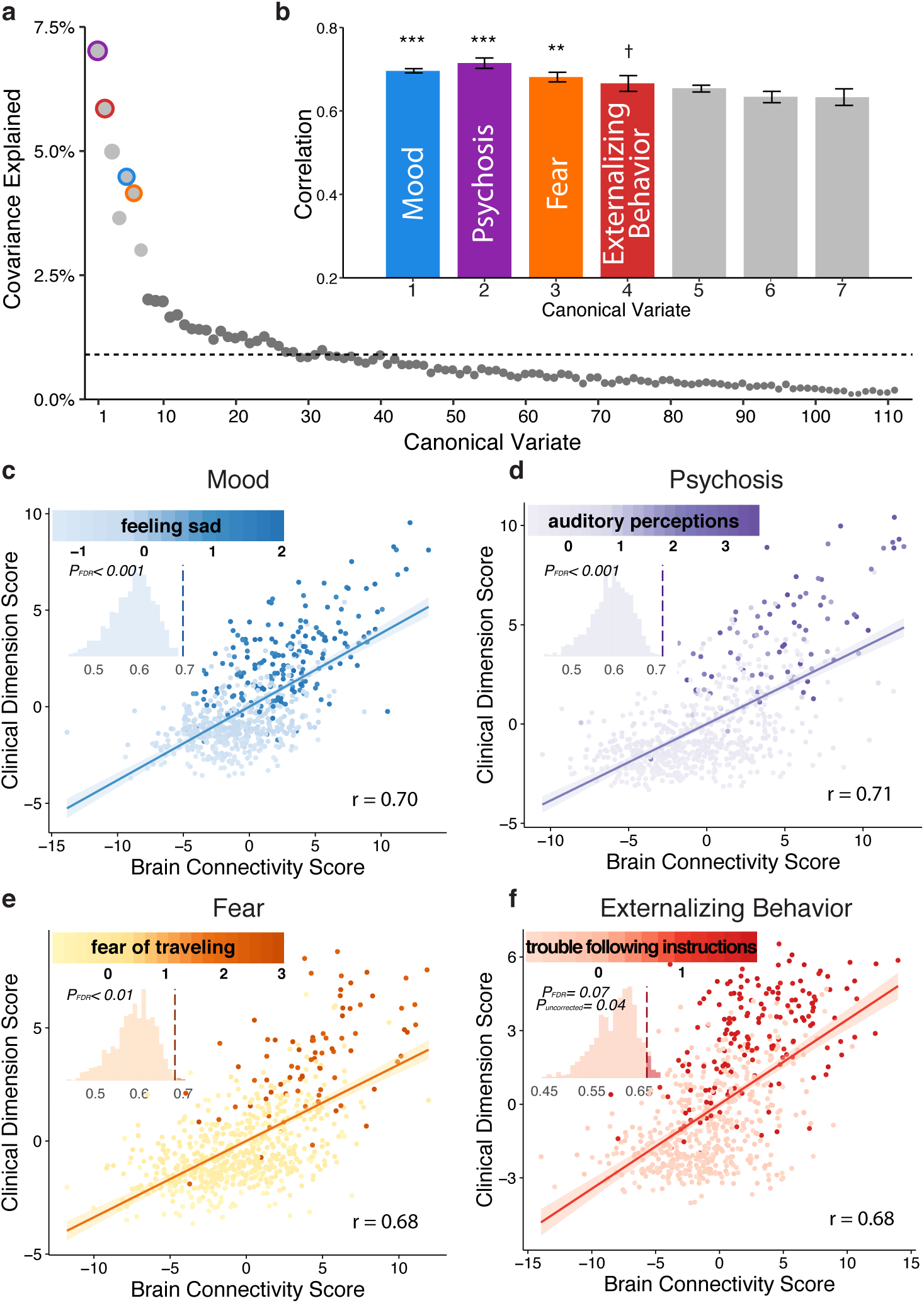
sCCA reveals multivariate patterns of linked dimensions of psychopathology and connectivity. **(a)** The first seven canonical variates were selected based on covariance explained. Dashed line marks the average covariance explained. **(b)** Three canonical correlations were statistically significant by permutation testing with FDR correction (*q <* 0.05), with the fourth one showing an effect at uncorrected thresholds. Corresponding variates are circled in (a). Error bars denote standard error. Dimensions are ordered by their permutation-based P value. **(c-f)** Scatter plots of brain and clinical scores (linearcombinations of functional connectivity and psychiatric symptoms, respectively) demonstrate the correlated multivariate patterns of connectomic and clinical features. Colored dots in each panel indicate the severity of a representative clinical symptom that contributed the most to this canonical variate. Each insert displays the null distribution of sCCA correlation by permutation testing. Dashed line marks the actual correlation. \*\*\**P*_*FDR*_ < 0.001, \*\**P*_*FDR*_ < 0.01, *†Puncorrected* = 0.04.

Each canonical variate represented a distinct pattern that relates a weighted set of psychiatric symptoms to a weighted set of functional connections. Inspection of the most heavily weighted clinical symptom for each dimension provided an initial indication regarding their content (**Fig. 2c-f**). For example, “feeling” sad was the most heavily weighted clinical feature in the first dimension, while “auditory perceptions” was the most prominent symptom in the second. Next, we conducted detailed analyses to describe the clinical and connectivity features driving the observed multivariate relationships.

### Interpretable, connectivity-guided dimensions of psychopathology cross clinical diagnostic categories

To understand the characteristics of each linked dimension, we used a resampling procedure to identify both clinical and connectivity features that were consistently significant across subsets of the data (Online Methods and see **Supplementary Fig. 6)**. This procedure revealed that 37 out of 111 psychiatric symptoms reliably contributed to at least one of the four dimensions (**Fig. 3)**. Next, we mapped these data-driven items to typical clinical diagnostic categories. This revealed that the features selected by multivariate analyses generally accord with clinical phenomenology. Specifically, despite being selected on the basis of their relationship with functional connectivity, the first three canonical variates delineated dimensions that resemble clinically coherent dimensions of mood, psychosis, and fear (e.g. phobias). The fourth dimension, which was present at an uncorrected threshold, mapped to externalizing behaviors (attention deficit/hyperactivity disorder (ADHD) and oppositional defiant disorder (ODD)).

**Figure 3.**
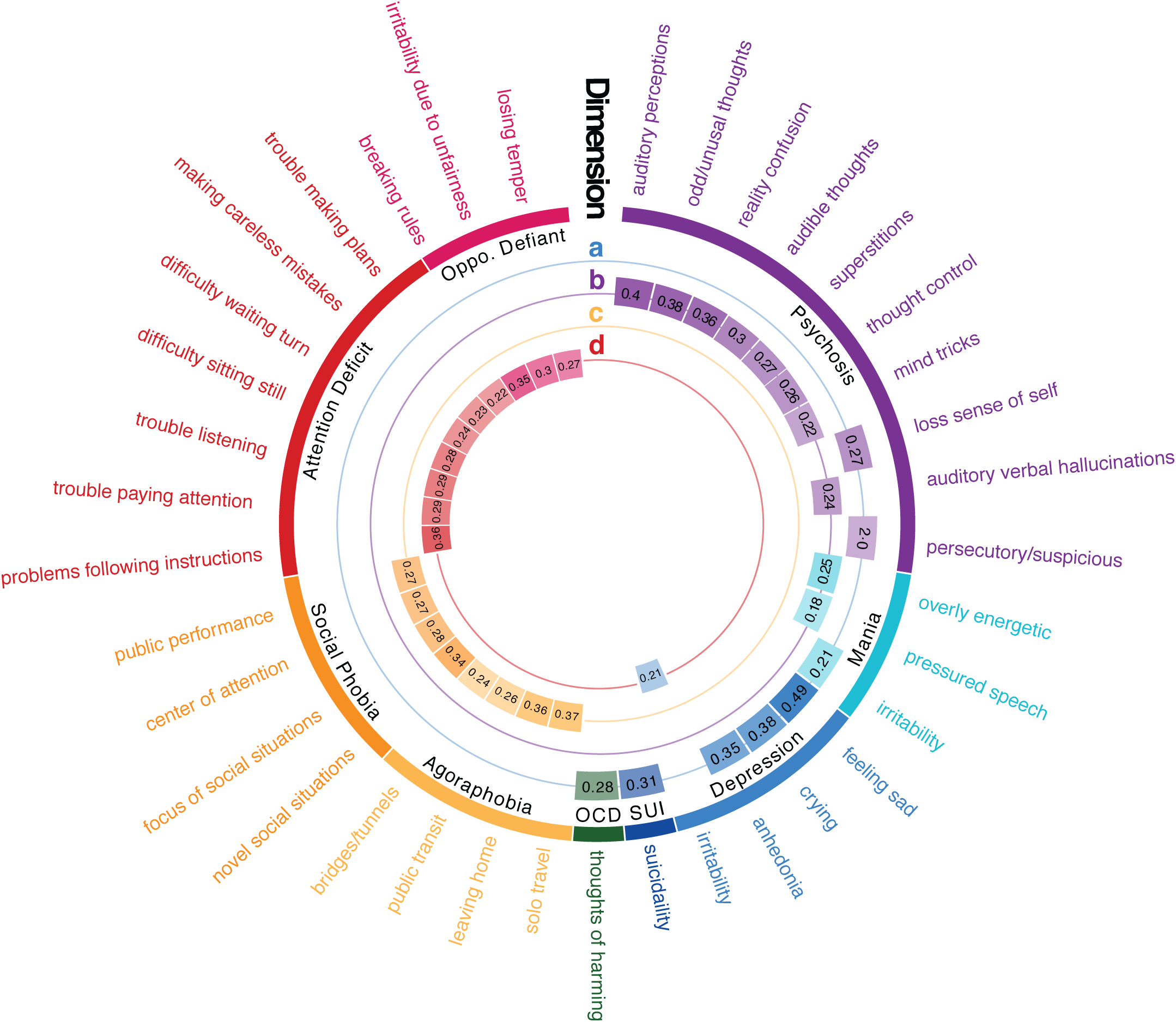
Connectivity-informed dimensions of psychopathology cross clinical diagnostic categories. **(a)** The mood dimension was composed of a mixture of depressive symptoms, suicidality, irritability, and recurrent thoughts of self harm. **(b)** The psychotic dimension was composed of psychosis-spectrum symptoms, as well as two manic symptoms. **(c)** The fear dimension was comprised of social phobia and agoraphobia symptoms. **(d)** The externalizing behavior dimension showed a mixture of symptoms from attention-deficit and oppositional defiant disorders, as well as irritability from the depression section. The outermost labels are the item-level psychiatric symptoms. The color arcs represent categories from clinical screening interview and the Diagnostic and Statistical Manual of Mental Disorders (DSM). Numbers in the inner rings represent sCCA loadings for each symptom in their respective dimension. Only loadings determined to be statistically significant by a resampling procedure are shown here.

While each canonical variate mapped onto coherent clinical features, each dimension contained symptoms from several different clinical diagnostic categories. For example, the mood dimension was comprised of symptoms from categorical domains of depression (“feeling sad” received the highest loading), mania (“irritability”), and obsessive-compulsive disorder (OCD; “recurrent thoughts of harming self or others”) (**Fig. 3a**). Similarly, while the second dimension mostly consisted of psychosis-spectrum symptoms (such as “auditory verbal hallucinations”), two manic symptoms (i.e. “overly energetic” and “pressured speech”) were included as well (**Fig. 3b**). The third dimension was composed of fear symptoms, including both agoraphobia and social phobia (**Fig. 3c**). The fourth dimension was driven primarily by symptoms of both ADHD and ODD, but also included the irritability item from the depression domain (**Fig. 3d**). These data-driven dimensions of psychopathology align with clinical phenomenology, but in a dimensional fashion that does not adhere to discrete categories.

### Common and dissociable patterns of dysconnectivity contribute to linked psychopathological dimensions

sCCA identified each dimension of psychopathology through shared associations between clinical data and specific patterns of dysconnectivity. Next, we investigated the loadings of connectivity features that underlie each canonical variate. To aid visualization of the high-dimensional connectivity data, we summarized loading patterns according to network communities established *a priori* by the parcellation system. Specifically, we examined patterns of both *within*-network and *between*-network connectivity (**Supplementary Fig. 7;** Online Methods), as this framework was useful in prior investigations of both brain development^46,52^ and psychopathology.^53–55^ This procedure revealed that the mood dimension was associated with increased connectivity *within* three networks: default mode, fronto-parietal, and salience networks (**Fig. 4a,e,i**). However, the most heavily weighted features in the mood dimension reflected abnormalities of connectivity *between* networks. In particular, mood was associated with a lack of segregation between the default mode and both the fronto-parietal and salience networks. The psychosis dimension similarly showed elevated connectivity within the default mode network and its reduced segregation from executive networks (fronto-parietal and salience)(**Fig. 4b,f,j**). The fear dimension also showed elevated connectivity within the fronto-parietal and salience networks, but in contrast showed reduced connectivity within the default mode network itself (**Fig. 4c,g,k**). As was the case for mood and psychosis, the fear dimension exhibited a failure of segregation between the default mode and executive networks (**Fig. 4c,g,k**). Reduced connectivity within the default mode network was also present in the externalizing dimension, as was reduced segregation between default mode and executive networks (**Fig. 4d,h,l**).

**Figure 4.**
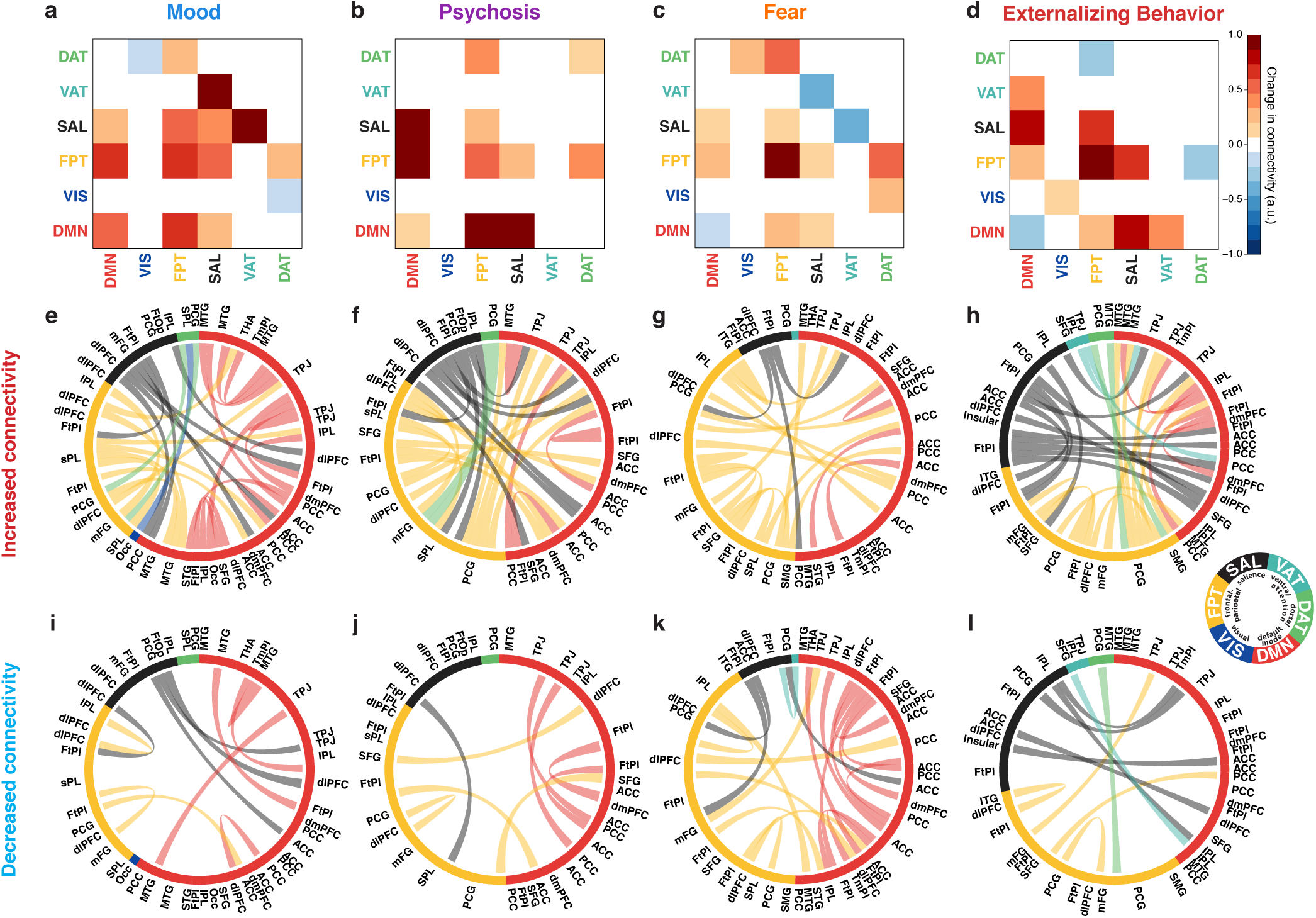
Specific patterns of within- and between-network dysconnectivity contribute to linked psychopathological dimensions. **(a-d)** Modular (community) level connectivity pattern associated with each psychopathology dimension. Both increased **(e-h)** and diminished **(i-l)** connectivity in specific edges contributed to each dimension of psychopathology. The outer labels represent the anatomical names of nodes. The inner arcs indicate the community membership of nodes. The thickness of the chords represent the loadings of connectivity features.

The results indicate that while each canonical variate was comprised of unique patterns of dysconnectivity, there were several features that were shared across all dimensions. Such findings agree with accumulating evidence for common circuit-level dysfunction across psychiatric syndromes.^6–10^ To quantitatively assess such common features, we compared overlapping results against a null distribution using permutation testing (see Online Methods). This procedure revealed an ensemble of edges that were consistently implicated across all four dimensions. These connections can be mapped to individual nodes, and revealed that the regions most impacted across all dimensions included the frontal pole, superior frontal gyrus, dorsomedial prefrontal cortex, medial temporal gyrus, and amygdala (**Fig. 5a**). Similar analysis at the level of sub-networks (**Fig. 5b**) illustrated that abnormalities of connectivity *within* the default mode and fronto-parietal networks were present in all four psychopathological dimensions (**Fig. 5c**). Furthermore, reduced segregation *between* the default mode and executive networks, such as the fronto-parietal and salience systems, was common to all dimensions. These shared connectivity features complement each dimension-specific pattern, and offer evidence for both common and dissociable patterns of dysconnectivity associated with psychopathology.

**Figure 5.**
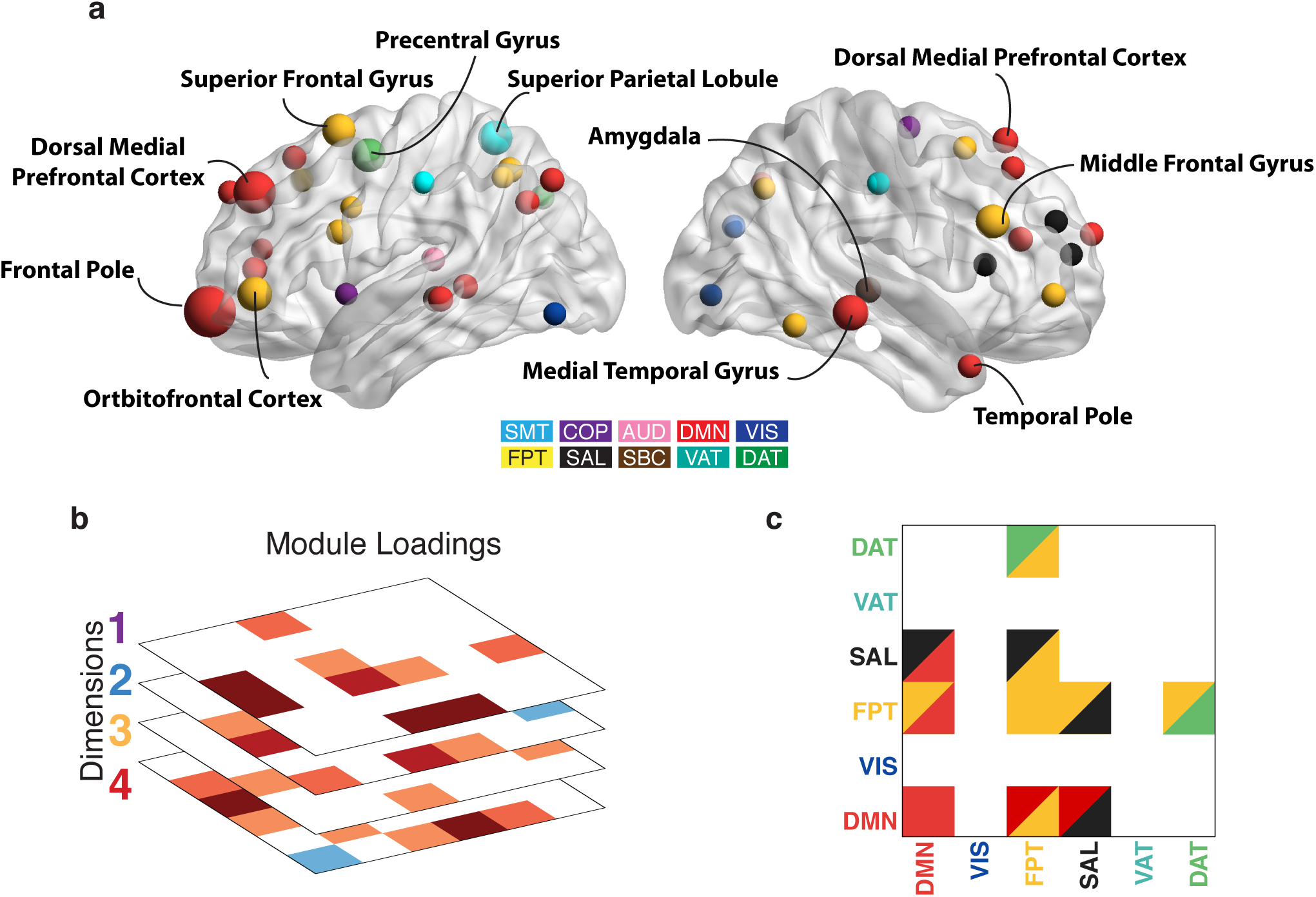
Loss of segregation between default mode and executive networks is shared across all dimensions. **(a)** By searching for overlap of edges that contributed significantly to each dimension, we found common edges that were implicated across all dimensions of psychopathology. These were then summarized at a nodal level by the sum of their absolute loadings. Nodes that contributed significantly to every dimension included the frontal pole, superior frontal gyrus, dorsomedial prefrontal cortex, medial temporal gyrus, and amygdala. **(b)** Results of a similar analysis conducted at the module level. **(c)** Loss of segregation between the default mode and executive networks were shared across all four dimensions.

### Developmental effects and sex differences are concentrated in specific dimensions

In the above analyses, we examined multivariate associations between connectivity and psychopathology while controlling for participant age. However, given that abnormal neurodevelopment is thought to underlie many psychiatric disorders,^41–44^ we next examined whether connectivity patterns significantly associated with psychopathology differ as a function of age or sex in this large developmental cohort. We repeated the analysis conducted above using connectivity and clinical features, but in this case did not regress out age and sex; race and motion were still regressed from both datasets. Notably, the dimensions derived were quite similar, with highly correlated feature weights (**Supplementary Table 2**). As in prior work,^40,56–58^ developmental associations were examined using generalized additive models with penalized splines, which allows for statistically rigorous modeling of both linear and non-linear effects while minimizing over-fitting. Using this approach, we found that the brain connectivity patterns associated with both mood and psychosis became significantly more prominent with age (**Fig. 6a,b**, *P*_*FDR*_ < 10^−13^, *P*_*FDR*_ < 10^−6^, respectively). Additionally, brain connectivity patterns linked to mood and fear were both stronger in female participants than males (**Fig. 6c,d**, *P*_*FDR*_ < 10^−8^, *P*_*FDR*_ < 10^−7^, respectively). We did not observe age by sex interaction effects in any dimension.

**Figure 6.**
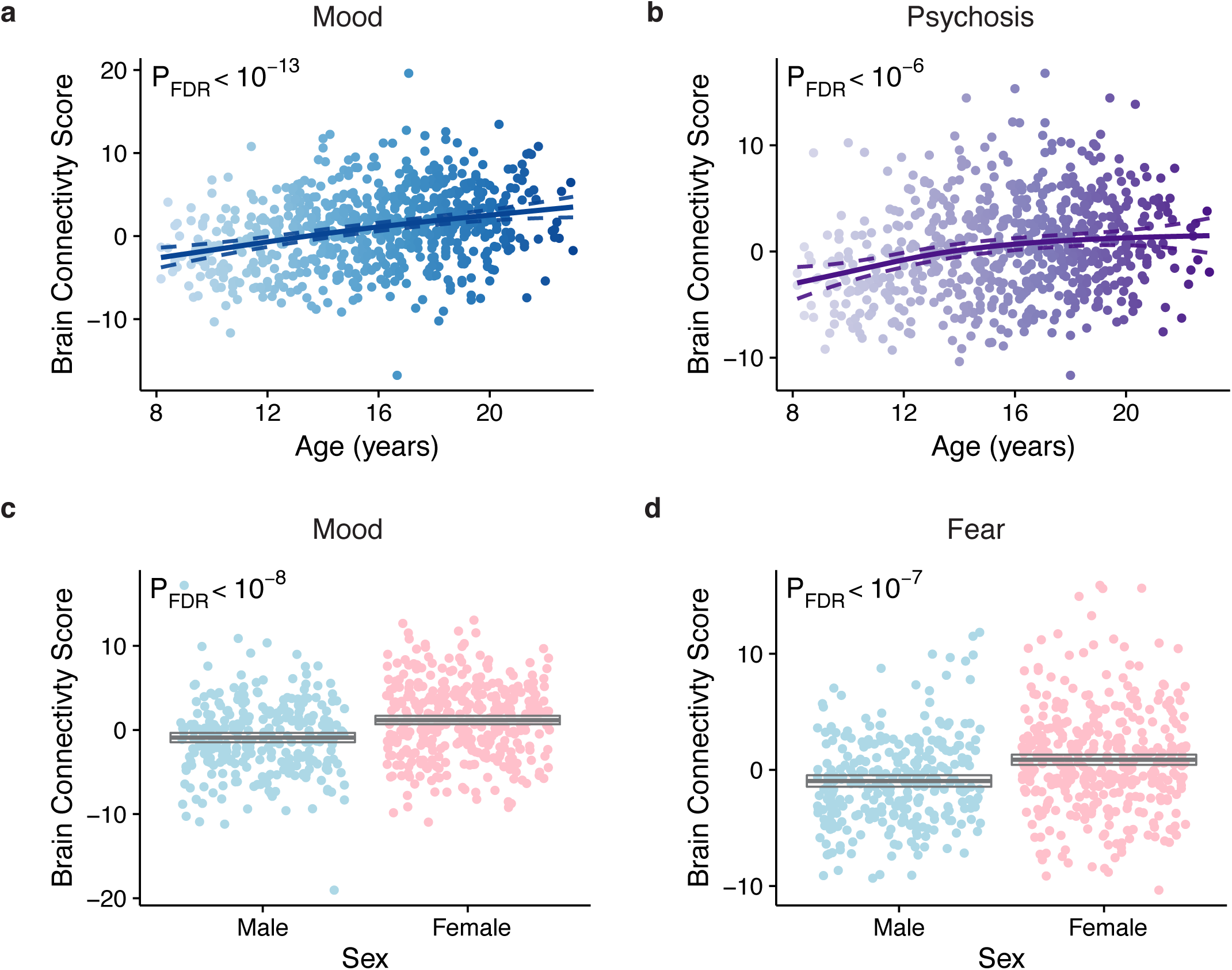
Developmental effects and sex differences are concentrated in specific dimensions. Connectivity patterns associated with both the mood **(a)** and psychosis **(b)** dimensions increased significantly with age. Additionally, connectivity patterns associated with both the mood **(c)** and fear **(d)** symptoms were significantly more prominent in females than males. Multiple comparisons were controlled for using the False Discovery Rate (*q <* 0.05).

### Linked dimensions are replicated in an independent sample

Throughout our analysis of the discovery sample, we used procedures both to guard against over-fitting and to enhance the generalizablity of results (regularization, permutation testing, resampling). As a final step, we tested the replicability of our findings using an independent sample, which was left-out from all analyses described above (n=336, **Supplementary Fig. 1)**. Although this replication sample was half the size of our original discovery sample, sCCA identified four canonical variates that highly resemble the original four linked dimensions of psychopathology (with correlations of loadings between discovery and replication within 0.24 and 0.88; **Fig. 7a,b**). In the replication sample, three out of four dimensions were significant after FDR correction of permutation tests (**Supplementary Fig. 8)**.

**Figure 7.**
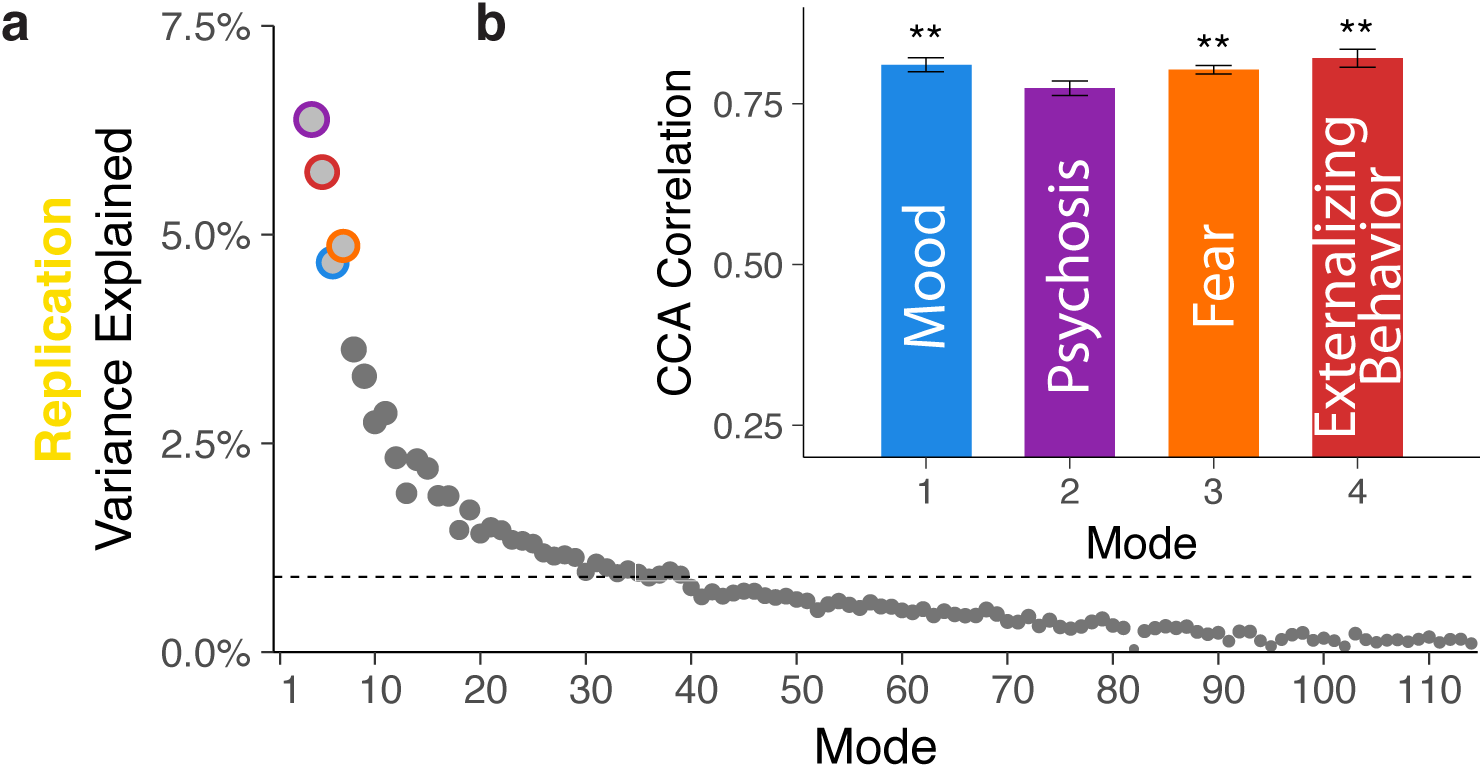
Linked dimensions of psychopathology were replicated in an independent sample. All procedures were repeated in an independent replication sample of 336 participants. **(a)** The first four canonical variates in the replication sample were selected for further analysis based on covariance explained. Dashed line marks the average covariance explained. **(b)** The mood, fear, and externalizing behavior dimensions were significant by permutation testing. Corresponding variates are circled in (a). Error bars denote standard error. \*\**P*_*FDR*_ < 0.01.

## DISCUSSION

Leveraging a large neuroimaging data set of youth and recent advances in machine learning, we discovered several multivariate patterns of functional dysconnectivity linked to interpretable dimensions of psychopathology that cross traditional diagnostic categories. These patterns of abnormal connectivity were both highly correlated and replicable in an independent dataset. While each dimension displayed a specific pattern of connectivity abnormalities, loss of network segregation between the default mode and executive networks was common to all dimensions. Furthermore, patterns of dysconnectivity displayed unique developmental effects and sex differences. Together, these results suggest that complex psychiatric symptoms are associated with specific patterns of abnormal connectivity during brain development.

Both the co-morbidity among psychiatric diagnoses and the notable heterogeneity within each diagnostic category suggest that our current symptom-based diagnostic criteria do not ″carve nature at its joints″.^2–4^ Establishing biologically-targeted interventions in psychiatry is predicated upon delineation of the underlying neurobiology. This challenge has motivated the NIMH Research Domain Criteria (RDoC) effort, which seeks to link circuit-level abnormalities in specific brain systems to symptoms that might be present across clinical diagnoses.^11,12^ Accordingly, there has been a proliferation of studies that focus on linking specific brain circuit(s) to a specific symptom dimension or behavioral measure across diagnostic categories.^9,10, 29, 56, 59–63^ However, by focusing on a single behavioral measure or symptom domain, many studies ignore the co-morbidity among psychiatric symptoms. A common way to attempt to evaluate such co-morbidity is to find latent dimensions of psychopathology using factor analysis or related techniques.^57,64, 65^ For example, factor analyses of clinical psychopathology have suggested the presence of dimensions including internalizing symptoms, externalizing symptoms, and psychosis symptoms.^50^ While such dimensions are reliable, they are drawn entirely from the covariance structure of self-report or interview-based clinical data, and are not informed by neurobiology.

An alternative increasingly pursued is to parse heterogeneity in psychiatric conditions using multivariate analysis of biomarker data such as neuroimaging. For example, researchers have used functional connectivity^59^ and gray matter density^66^ to study the heterogeneity within major depressive disorder and psychotic disorders, respectively. However, most studies have principally considered only one or two clinical diagnostic categories, and typically the analytic approach yields discrete subtypes (or “biotypes”). By definition, such a design is unable to discover continuous dimensions that span multiple categories. Further, there is tension between the dimensional schema suggested by RDoC and categorical biotypes; as suggested by RDoC, it seems more plausible that psychopathology in an individual results from a mixture of abnormalities across several brain systems. Finally, unsupervised learning approaches using only imaging data and not considering clinical data may frequently yield solutions that are difficult to interpret, and that do not align with clinical experience.

In contrast, in this study we used a multivariate analysis technique – sCCA – that allowed simultaneous consideration of clinical and functional connectivity data in a large sample with diverse psychopathology. This enabled uncovering linked dimensions of psychopathology and dysconnectivity that cross diagnostic categories yet remain clinically interpretable. In contrast to “one-view” multivariate studies^50,57, 64, 67^ (such asfactor analysis of clinical data or clustering of imaging data), the sCCA-derived clinical dimensions were explicitly selected on the basis of co-varying signals that were present as both alterations of connectivity and clinical symptoms. Such a “two-view” approach has been successfully applied in studies of neurodegenerative diseases^48^ and normal brain-behavior relationships.^49^

Notably, the brain-driven dimensions described here incorporated symptoms across several diagnostic categories while remaining congruent with prevailing models of psychopathology. For example, the mood dimension was composed of items from five sections of the clinical interview: depression, mania, OCD, suicidality, and psychosis-spectrum. Despite disparate origins, the content of the items forms a clinically coherent picture, including depressed mood, anhedonia, loss of sense of self, recurrent thoughts of self harm, and irritability. Notably, symptoms of irritability were also significantly represented in the externalizing behavior dimension, suggesting that irritability may have heterogeneous, divergent neurobiological antecedents. The fear dimension, on the other hand, represents a more homogeneous picture of various types of phobias (e.g. social phobia and agoraphobia), that had little overlap with other categorical symptoms. Finally, the psychosis dimension (which was only significant in the discovery sample) was mainly comprised of psychotic symptoms, but also included symptoms of mania. This result accords with studies demonstrating shared inheritance patterns of schizophrenia and bipolar disorder, and findings that specific common genetic variants increase risk of both disorders.^68^ Instead of averaging over many clinical features within a diagnostic category, sCCA selected specific items that are most tightly linked to patterns of dysconnectivity. These groups of symptoms remained highly interpretable, and were reproducible in the replication data set.

Each of the clinical dimensions identified was highly correlated with patterns of dysconnectivity. These patterns were summarized according to their location *between* and *within* functional network modules, which has been a useful framework for understanding both brain development and psychopathology.^53–55^ While each dimension of psychopathology was associated with a unique pattern of dysconnectivity, one of the most striking findings to emerge was evidence that reduction of functional segregation between the default mode and fronto-parietal networks was a common feature of all dimensions. The exact connections implicated in each dimension might vary, but permutation-based analyses demonstrated that loss of segregation between these two networks was present in all four dimensions. Fox et al.^69^ originally demonstrated that the default mode network is anti-correlated with task-positive functional brain systems including the fronto-parietal network. Furthermore, studies of brain maturation have shown that age-related segregation of functional brain modules is a robust and reproducible findings regarding adolescent brain development.^38–40^ As part of this process, connections within network modules strengthen and connections between two network modules weaken. This process is apparent using functional connectivity^38,39^ as well as structural connectivity.^40^ Notably, case-control studies of psychiatric disorders in adults have found abnormalities consistent with a failure of developmental network segregation, in particular between executive networks, such as the fronto-parietal and salience networks, and the default mode network.^27,29^ Using a purely data-driven analysis, our results support the possibility that loss of segregation between the default mode and executive networks may be a common neurobiological mechanism underlying vulnerability to a wide range of psychiatric symptoms.

In addition to such common abnormalities that were present across dimensions, each dimension of psychopathology was associated with a unique, highly correlated pattern of dysconnectivity. For example, connectivity features linked to the mood dimension included hyper-connectivity within the default mode, fronto-parietal and salience networks. These dimensional results from a multivariate analysis are remarkably consistent with prior work, which has provided evidence of default mode hyper-connectivity using conventional case-control designs and univariate analysis.^55,70–73^ However, the data-driven approach used here allowed us to discover a combination of novel connectivity features that was more predictive than traditional univariate association analyses. These features included enhanced connectivity between both the dorsal attention and fronto-parietal networks as well as between the ventral attention and salience networks. The fear, externalizing, and psychosis dimensions were defined by a similar mix between novel features and a convergence with prior studies. Specifically, fear was characterized by weakened connectivity within default mode network, enhanced connectivity within fronto-parietal network, and — in contrast to mood — decreased connectivity between ventral attention and salience networks. In contrast to other dimensions, externalizing behavior exhibited increased connectivity in the visual network and decreased connectivity between fronto-parietal and dorsal attention networks.

Importantly, each of these dimensions was initially discovered while controlling for the effects of age and sex. However, given that many psychiatric symptoms during adolescence show a clear evolution with development and marked disparities between males and females,^44,74^ we evaluated how the connectivity features associated with each dimension were correlated with age and sex. We found that the patterns of dysconnectivity that linked to mood and psychosis symptoms strengthened with age during the adolescent period. This finding is consistent with the well-described clinical trajectory of both mood and psychosis disorders, which often emerge in adolescence and escalate in severity during the transition to adulthood.^75,76^ In contrast, no age effects were found for externalizing or fear symptoms, which are typically present earlier in childhood and have a more stable time-course.^77,78^ Additionally, we observed marked sex differences in the patterns of connectivity that linked to mood and fear symptoms, with these patterns being more prominent in females across the age range studied. This result accords with data from large-scale epidemiological studies, which have documented a far higher risk of mood and anxiety disorders in females.^79,80^ Despite marked differences in risk by sex (i.e. double in some samples), the mechanism of such vulnerability has been only sparsely studied in the past.^46,56, 62^ The present results suggest that sex differences in functional connectivity may in part mediate the risk of mood and fear symptoms.

Although this study benefited from a large sample, advanced multivariate methods, and replication of results in an independent sample, several limitations should be noted. First, although the item-level data used do not explicitly consider clinical diagnostic categories, the items themselves were nonetheless drawn from a standard clinical interview. Incorporating additional data types such as genomics may capture different sources of important biological heterogeneity. Second, while we successfully replicated our findings (except for the psychosis dimension) in an independent sample, the generalizability of the study should be further evaluated in datasets that are acquired in different settings. Third, all data considered in this study were cross-sectional, which has inherent limitations for studies of development. Ongoing follow-up of this cohort will yield informative data that will allow us to evaluate the suitability of these brain-derived dimensions of psychopathology for charting developmental trajectories and prediction of clinical outcome.

In summary, in this study we discovered and replicated multivariate patterns of connectivity that are highly correlated with dimensions of psychopathology in a large sample of youth. These dimensions cross traditional clinical diagnostic categories, yet align with clinical experience. Each dimension was composed of unique features of dysconnectivity, while a lack of functional segregation between the default mode network and executive networks was common to all dimensions. Paralleling the clinical trajectory of each disorder and known disparities in prevalence between males and females, we observed both marked developmental effects and sex differences in these patterns of dysconnectivity. As suggested by the NIMH Research Domain Criteria, our findings demonstrate how specific circuit-level abnormalities in the brain’s functional network architecture may give rise to a diverse panoply of psychiatric symptoms. Such an approach has the potential to clarify the high co-morbidity between psychiatric diagnoses and the great heterogeneity within each diagnostic category. Moving forward, the ability of these dimensions to predict disease trajectory and response to treatment should be evaluated, as such a neurobiologically-grounded framework could accelerate the rise of personalized medicine in psychiatry.

## End Notes

### Acknowledgements

Thanks to Chad Jackson for data management and systems support. This study was supported by grants from National Institute of Mental Health: R01MH107703 (TDS), R01MH112847 (RTS TDS), R21MH106799 (DSB TDS), R01MH107235 (RCG), and R01EB022573 (CD). The PNC was supported by MH089983 and MH089924. Additional support was provided by P50MH096891 to REG, R01MH101111 to DHW, K01MH102609 to DRR, K08MH079364 to MEC, R01NS085211 to RTS, and the Dowshen Program for Neuroscience. Data deposition: The data reported in this paper have been deposited in database of Genotypes and Phenotypes (dbGaP), www.ncbi.nlm.nih.gov/gap (accession no. phs000607.v1.p1). All analysis code is available here https://github.com/cedricx/sCCA/tree/master/sCCA/code/final.

### Author Contributions

All authors contributed to this work, approved the final manuscript, and agreed to the copyright transfer policies of the journal.

## Online Methods

### Participants

Resting-state functional magnetic resonance imaging (rs-fMRI) datasets were acquired as part of the Philadelphia Neurodevelopmental Cohort (PNC), a large community-based study of brain development.^46^ In total, 1601 participants completed the cross-sectional neuroimaging protocol (**Supplementary Fig. 1a**). One subject had missing clinical data. To create two independent samples for discovery and replication analyses, we performed random split of the remaining 1600 participants using the caret package^82^ in R. Specifically, using the function CreateDataPartition, a discovery sample (n=1069) and a replication sample (n=531) were created that were stratified by overall psychopathology. The two samples were confirmed to also have similar distributions in regards to age, sex, and race (**Supplementary Fig. 1b**). The overall psychopathology is the general factor score reported previously from factor analysis of the clinical data alone.^50,57^

Of the discovery sample (n=1069), 111 were excluded due to: gross radiological abnormalities, or a history of medical problems that might affect brain function. Of the remaining 958 participants, 45 were excluded for having low quality T1-weighted images, and 250 were excluded for missing rs-fMRI, incomplete voxelwise coverage, or excessive motion during the functional scan, which is defined as having an average framewise motion more than 0.20mm or more than 20 frames exhibiting over 0.25mm movement. These exclusion criteria produced a final discovery sample consisting of 663 youths (mean age 15.82, SD = 3.32; 293 males and 370 females). Applying the same exclusion criteria to the replication sample produced 336 participants (mean age 15.65, SD = 3.32; 155 males and 181 females). See **Supplementary Table 1** for detailed demographics of each sample.

### Psychiatric assessment

Psychopathology symptoms were evaluated using a structured screening interview (GOASSESS), which has been described in detail elsewhere.^50^ To allow rapid training and standardization across a large number of assessors, GOASSESS was designed to be highly structured, with screen-level symptom and episode information. The instrument is abbreviated and modified from the epidemiologic version of the NIMH Genetic Epidemiology Research Branch Kiddie-SADS.^81^ The psychopathology screen in GOASSESS assessed lifetime occurrence of major domains of psychopathology including psychosis spectrum symptoms, mood (major depressive episode, mania), anxiety (agoraphobia, generalized anxiety, panic, specific phobia, social phobia, separation anxiety), behavioral disorders (oppositional defiant, attention deficit/hyperactivity, conduct) disorders, eating disorders (anorexia, bulimia), and suicidal thinking and behavior. The 111 item-level symptoms used in this study were described in prior factor analysis of the clinical data in PNC.^57^ For the specific items, see **Supplementary Table 3**.

### Image acquisition

Structural and functional subject data were acquired on a 3T Siemens Tim Trio scanner with a 32-channel head coil (Erlangen, Germany), as previously described.^46,62^ High-resolution structural images were acquired in order to facilitate alignment of individual subject images into a common space. Structural images were acquired using a magnetization-prepared, rapid-acquisition gradient-echo (MPRAGE) T1-weighted sequence (*T*_*R*_ = 1810ms; *T*_*E*_ = 3.51ms; FoV = 180 *×* 240mm; resolution 0.9375 *×* 0.9375 *×* 1mm). Approximately 6 minutes of task-free functional data were acquired for each subject using a blood oxygen level-dependent (BOLD-weighted) sequence (*T*_*R*_ = 3000ms; *T*_*E*_ = 32ms; FoV = 192 *×* 192mm; resolution 3mm isotropic; 124 volumes). Prior to scanning, in order to acclimate subjects to the MRI environment and to help subjects learn to remain still during the actual scanning session, a mock scanning session was conducted using a decommissioned MRI scanner and head coil. Mock scanning was accompanied by acoustic recordings of the noise produced by gradient coils for each scanning pulse sequence. During these sessions, feedback regarding head movement was provided using the MoTrack motion tracking system (Psychology Software Tools, Inc, Sharpsburg, PA). Motion feedback was only given during the mock scanning session. In order to further minimize motion, prior to data acquisition subjects’ heads were stabilized in the head coil using one foam pad over each ear and a third over the top of the head. During the resting-state scan, a fixation cross was displayed as images were acquired. Subjects were instructed to stay awake, keep their eyes open, fixate on the displayed crosshair, and remain still.

### Structural Preprocessing

A study-specific template was generated from a sample of 120 PNC subjects balanced across sex, race, and age bins using the buildTemplateParallel procedure in ANTS.^83^ Study-specific tissue priors were created using a multi-atlas segmentation procedure.^84^ Subject anatomical images were independently rated by three highly trained image analysts. Any image that did not pass manual inspection was removed from the analysis. Each subject’s high-resolution structural image was processed using the ANTS Cortical Thickness Pipeline.^85^ Following bias field correction,^86^ each structural image was diffeomorphically registered to the study-specific PNC template using the top-performing SYN deformation provided by ANTS.^87^ Study-specific tissue priors were used to guide brain extraction and segmentation of the subject’s structural image.^88^

### Functional Preprocessing

Task-free functional images were processed using one of the top-performing pipelines for removal of motion-related artifact.^51^ Preprocessing steps included (1) correction for distortions induced by magnetic field inhomogeneities using FSL’s FUGUE utility, (2) removal of the 4 initial volumes of each acquisition, (3) realignment of all volumes to a selected reference volume using MCFLIRT,^89^ (4) removal of and interpolation over intensity outliers in each voxel’s time series using AFNI’s 3DDESPIKE utility, (5) demeaning and removal of any linear or quadratic trends, and (6) co-registration of functional data to the high-resolution structural image using boundary-based registration.^90^ The artefactual variance in the data was modelled using a total of 36 parameters, including the 6 framewise estimates of motion, the mean signal extracted from eroded white matter and cerebrospinal fluid compartments, the mean signal extracted from the entire brain, the derivatives of each of these 9 parameters, and quadratic terms of each of the 9 parameters and their derivatives. Both the BOLD-weighted time series and the artefactual model time series were temporally filtered using a first-order Butterworth filter with a passband between 0.01 and 0.08 Hz.^91^

### Network construction

A functional connectivity network was computed across all parcels of a common parcellation using the residual timeseries following de-noising.^19^ The functional connectivity between any pair of brain regions was operationalised as the Pearson correlation coefficient between the mean activation timeseries extracted from those regions.^92^ For each parcellation, an *n × n* weighted adjacency matrix encoding the connectome was thus obtained, where *n* represents the total number of nodes (or parcels) in that parcellation. Community boundaries were defined *a priori* for each parcellation scheme. As part of the supplementary analysis to demonstrate the robustness of the results independent of parcellation choices (**Supplementary Fig. 5)**, we also constructed networks based on an alternative system.^24^

To ensure that potential confounders did not drive the canonical correlations, we regressed out relevant covariates out of the input matrices. Specifically, using the glm and residual.glm functions in R, we regressed age, sex, race and in-scanner motion out of the connectivity data, and regressed age, sex and race out of the clinical data. Importantly, we found that the canonical variates derived from regressed and non-regressed datasets were comparable, with highly correlated feature weights (**Supplementary Table 2**).

### Dimensionality reduction

Each correlation matrix comprised 34,980 unique connectivity features. We reasoned that since sCCA seeks to capture sources of variation common to both datasets, connectivity features that are most predictive of psychiatric symptoms would be those with high variance across participants. Therefore, to reduce dimensionality of the connectivity matrices, we selected the top edges with the highest median absolute deviation (MAD) (**Supplementary Fig. 2)**. MAD is defined as *median*(|**X***_i_ - median*(**X**)|), or the median of the absolute deviations from the vector’s median. We chose MAD as a measurement for variance estimation, because it is a robust statistic, being more resilient to outliers in a data set than other measures such as the standard deviation. To illustrate which edges were selected based on MAD, we visualized the network adjacency matrix with all edges, at 95th, 90th and 75th percentile (**Supplementary Fig. 2c**).

An alternative approach for dimensionality reduction is performing principal component analysis (PCA), from which we selected the top 111 components (explaining 37% of variance) as connectivity features entered into sCCA. As detailed in **Supplementary Fig. 5,** using PCA yielded similar canonical variates as MAD. We ultimately chose feature selection with MAD because it allowed direct use of individual connectivity strength instead of latent variables (e.g. components from PCA) as the input features to sCCA, thus increasing the interpretability of our results.

### sCCA

Sparse canonical correlation analysis (sCCA) is a multivariate procedure that seeks maximal correlations between linear combinations of variables in both sets,^93^ with regularization to achieve sparsity.^47^ In essence, given two matrices, **X***_n×p_* and **Y***_n×q_*, where *n* is the number of observations (e.g.participants), *p* and *q* are the number of variables (e.g. clinical and connectivity features, respectively), sCCA involves finding **u** and **v**, which are loading vectors, that maximize cor(**Xu**,**Yv**). Mathematically, this optimization problem can be expressed as

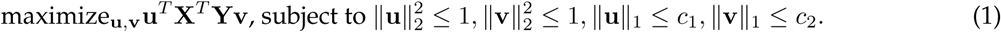

Since both *L*^1^ (∥x∥_1_) and *L*^2^-norm (∥x∥_2_) are used, this is an elastic net regularization that combines the LASSO and ridge penalties. The penalty parameters for the *L*^2^ norm are fixed for both u and v at 1, but those of *L*^1^ norm, namely *c*_1_ and *c*_2_, are set by the user and need to be tuned (see below).

### Grid search for regularization parameters

We tuned the *L*^1^ regularization parameters for the connectivity and the clinical features, respectively (see **Supplementary Fig. 3)**. The range of sparsity parameters are constrained to be between 0 and 1 in the PMA package,^47^ where 0 indicates the smallest number of features (i.e. highest level of sparsity) and 1 indicates the largest number of features (i.e. lowest level of sparsity). We conducted a grid search in increment of 0.1 to determine the combination of parameters that would yield the highest canonical correlation of the first variate across 10 randomly resampled samples, each consisting of two-thirds of the discovery dataset. Note that the parameters were only tuned on the discovery sample and the same regularization parameters were applied in the replication analysis.

### Permutation testing

To assess the statistical significance of each canonical variate, we used a permutation testing procedure to create a null distribution of correlations (**Supplementary Fig. 4)**. Essentially, we held the connectivity matrix constant, and then shuffled the rows of the clinical matrix so as to break the linkage of participants’ brain features and their symptom features. Then we performed sCCA using the same set of regularization parameters to generate a null distribution of correlations after permuting the input data 1000 times (B). As permutation could induce arbitrary axis rotation, which changes the order of canonical variates, or axis reflection, which causes a sign change for the weights, we matched the canonical variates resulting from permuted data matrices to the ones derived from the original data matrix by comparing the clinical loadings (v).^94^ The *P*_*FDR*_ value was estimated as the number of null correlations (*r*_*i*_) that exceeded the average sCCA correlations estimated on the original dataset 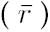 with false discovery rate correction (FDR, *q <* 0.05) across the top seven selected canonical variates:

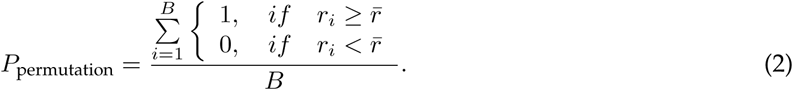

### Resampling procedure

To further select features that consistently contributed to each canonical variate, we performed a resampling procedure (**Supplementary Fig. 6)**. In each of 1000 samples, we randomly selected two-thirds of the discovery sample and then randomly replaced the remaining one-third from those two-thirds (similar to bootstrapping with replacement). Similar to the permutation procedure, we matched the corresponding canonical variates from resampled matrices to the original one to obtain a set of comparable decompositions.^94^ Features whose 95% and 99% confidence intervals (for clinical and connectivity features, respectively) did not cross zero were considered significant, suggesting that they were stable across different sampling cohorts.

### Network module analysis

To visualize and understand the high dimensional connectivity loading matrix, we summarized it as mean within- and between-module loadings according to the *a priori* community assignment of the Power^19^ parcellation (**Supplementary Fig. 7a**). Specifically, *within*-module connectivity loading is defined as

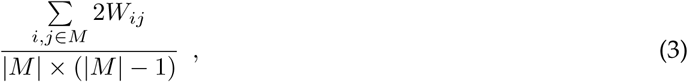

where *W*_*i,j*_ is the sCCA loading of the functional connectivity between nodes *i* and *j*, which both belong to the same community *m* in *M*. The cardinality of the community assignment vector, *|M|*, represents the number of nodes in each community. *Between*-module connectivity loading is defined as

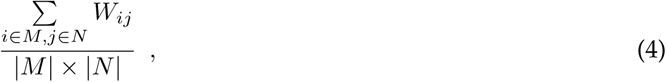

where *W*_*i,j*_ is the sCCA loading of the functional connectivity between nodes *i* and *j*, which belong to community *m* in *|M|* and community *n* in *|N|*, respectively.

We used a permutation test based on randomly assigning community memberships to each node while controlling for community size to assess the statistical significance of the mean connectivity loadings (**Supplementary Fig. 7b**). Empirical *P*-values were calculated similar to Equation 2 and were FDR corrected.

### Analysis of common connectivity features across dimensions

Each connectivity loading matrix was first binarized based on the presence of a significant edge feature after the resampling procedure in a given canonical variate. All four binarized matrices were then added and thresholded at 4 (i.e. common to all four dimensions), generating an overlapping edge matrix. Statistical significance was assessed by comparing this concordant feature matrix to a null model. The null model was constructed by computing the overlapping edges, repeated 1000 times, of four randomly generated loading matrices, each preserving the edge density of the original loading matrix. Any edge that appeared at least once in the null model was eliminated from further analysis. With only the statistically significant common edge features, we calculated the mean absolute loading in each edge feature across four dimensions as well as the nodal loading strength using `Brain Connectivity Toolbox`^95^ and visualized it with `BrainNet Viewer`^96^ both in MATLAB.

### Analysis of age effects and sex differences

As previously,^40,56–58^ generalized additive models (GAMs), using the MGCV package^97,98^ in R, were used to characterize age-related effects and sex differences on the specific dysconnectivity pattern associated with each psychopathology dimension. A GAM is similar to a generalized linear model where predictors can be replaced by smooth functions of themselves, offering efficient and flexible estimation of non-linear effects. For each linked dimension *i*, a GAM was fit:

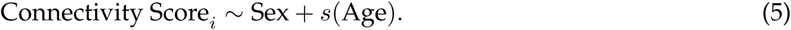

Additionally, we also separately tested whether age by sex interactions were present.

## Supplementary Information | Linked dimensions of psychopathology and connectivity in functional brain networks

**Supplementary Figure 1.**
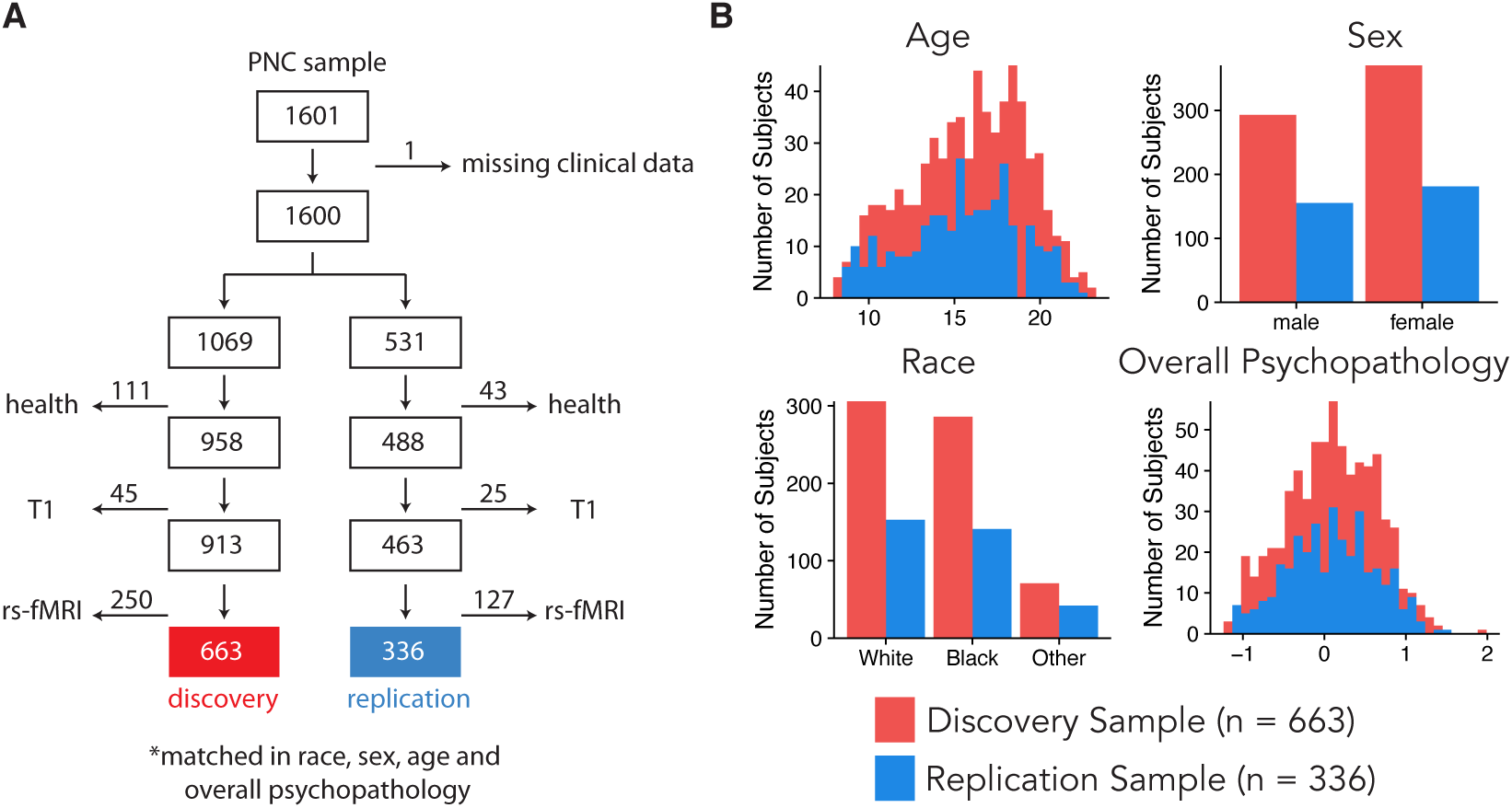
Sample Construction. **(a)** The cross-sectional sample of the Philadelphia Neurodevelopmental Cohort (PNC) has 1601 participants in total. Excluding the one missing clinical data, 1600 participants were randomly stratified into a discovery (n=1069) and a replication (n=531) sample. Applying health, structural and functional imaging quality exclusion criteria (details in Online Methods), 663 and 336 subjects were included in the final discovery and replication samples, respectively. **(b)** The two samples had similar demographic composition, including distributions of age, race, sex and overall psychopathology.

**Supplementary Figure 2.**
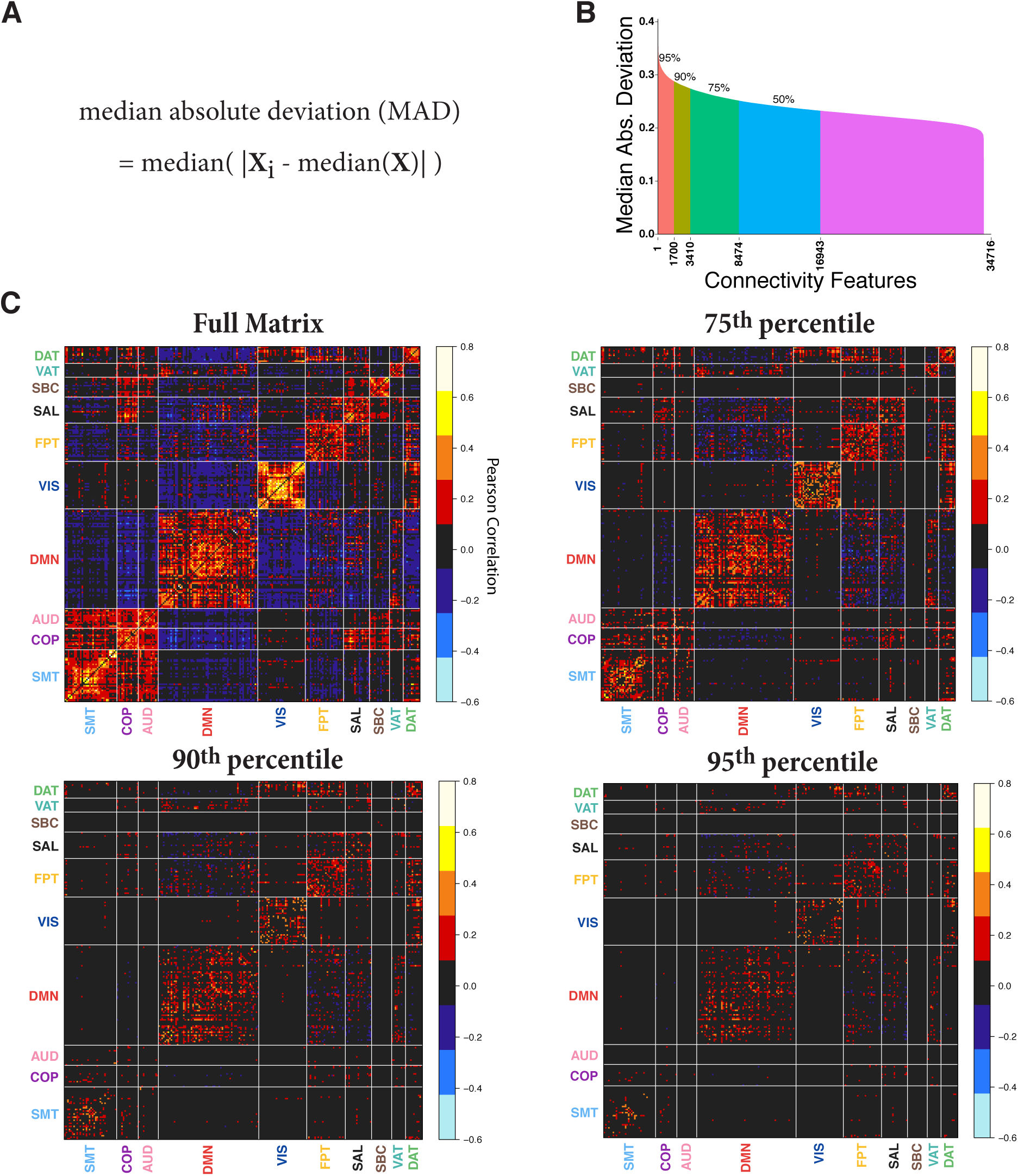
Connectivity feature selection using median absolute deviation (MAD). Since sCCA seeks to capture sources of variation common to both datasets, we selected top 10% or 3410 connectivity features that were variable across the discovery sample. **(a)** The variance was calculated using the median absolute deviation (MAD). It is defined as the median of the difference between each element and the median in a vector. **(b)** MAD of each edge strength in decreasing order. The 95th, 90th, and 75th percentile are labeled, where the 90th corresponds to 3410 edges. **(c)** Average connectivity matrix across all participants of edges with MAD at 100th, 95th, 90th, and 75th percentile levels.

**Supplementary Figure 3.**
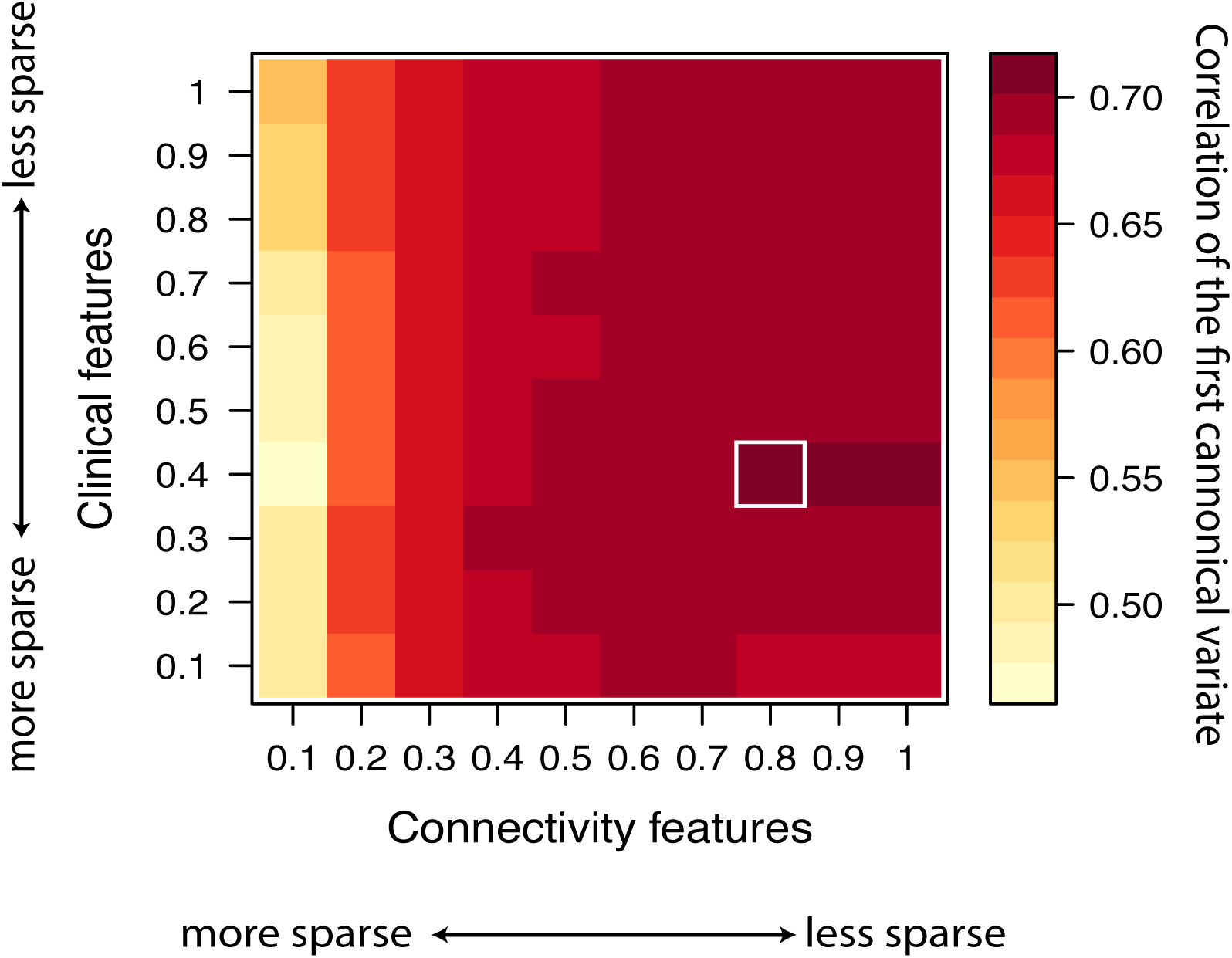
Grid search for regularization parameters. We tuned the *L*^1^ regularization parameters for the connectivity and the clinical features in sCCA. The range of sparsity parameters is constrained to be between 0 and 1 in the PMA package,^47^ where 0 indicates the smallest number of features (i.e. highest level of sparsity) and 1 indicates the largest number of features (i.e. lowest level of sparsity). We conducted a grid search in increment of 0.1 to determine the combination of parameters that would yield the highest canonical correlation of the first variate across 10 randomly resampled datasets, each consisting of two-thirds of the discovery dataset.

**Supplementary Figure 4.**
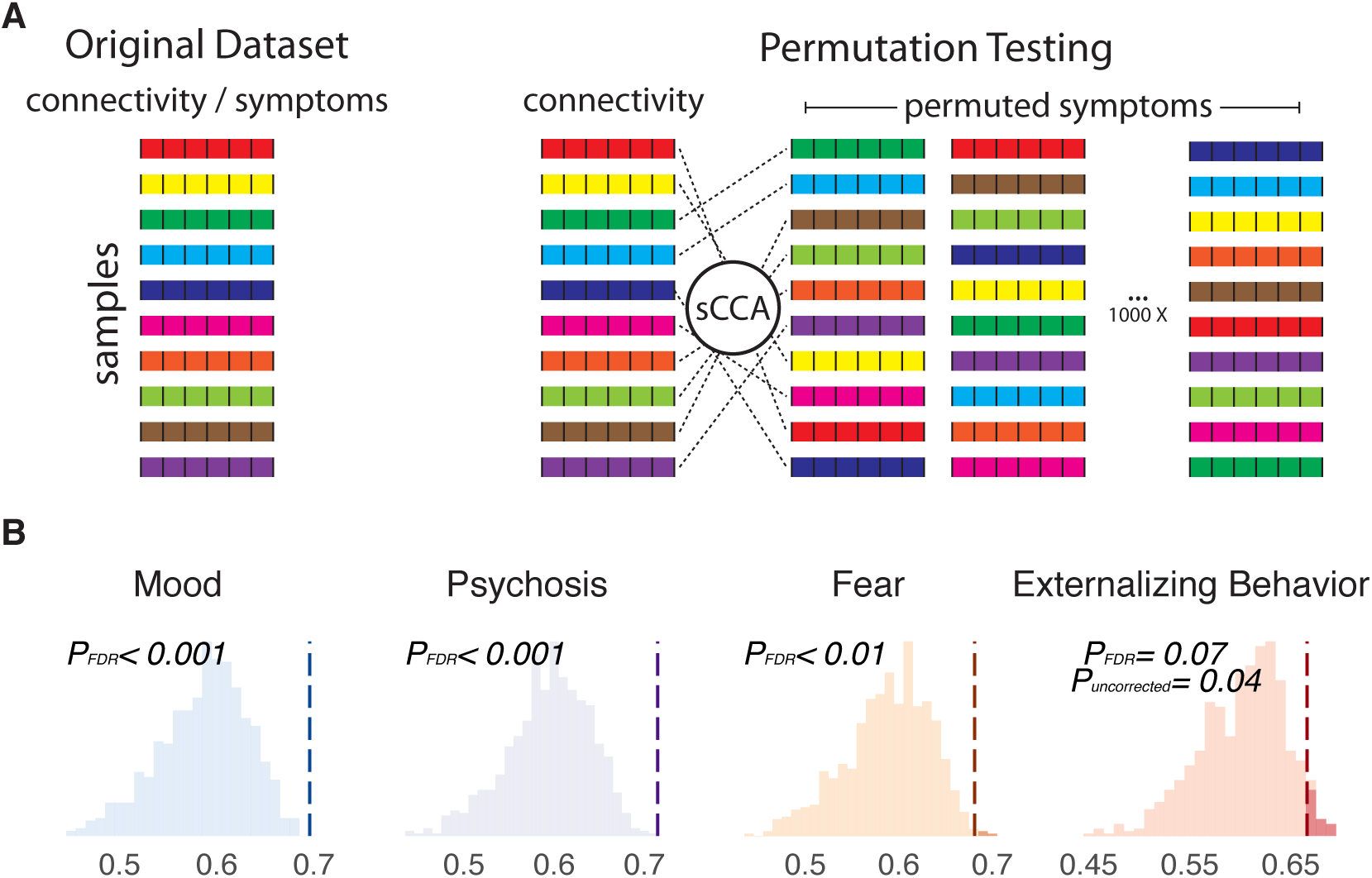
Permutation testing to assess significance of linked dimensions. **(a)** Schematic of permutation procedure. Connectivity data was held constant, while the rows of the clinical matrix were randomly shuffled, so as to break the linkage of participants’ connectivity features and their symptom features. As permutation could induce arbitrary axis rotation, which changes the order of canonical variates, or axis reflection, which causes a sign change for the weights, we matched the canonical variates resulting from permuted data matrices to the ones derived from the original data matrix by comparing the clinical loadings (v).^94^ **(b)** Null distributions of correlations generated by the permuted data. Dashed line represents the correlation from the original dataset.

**Supplementary Figure 5.**
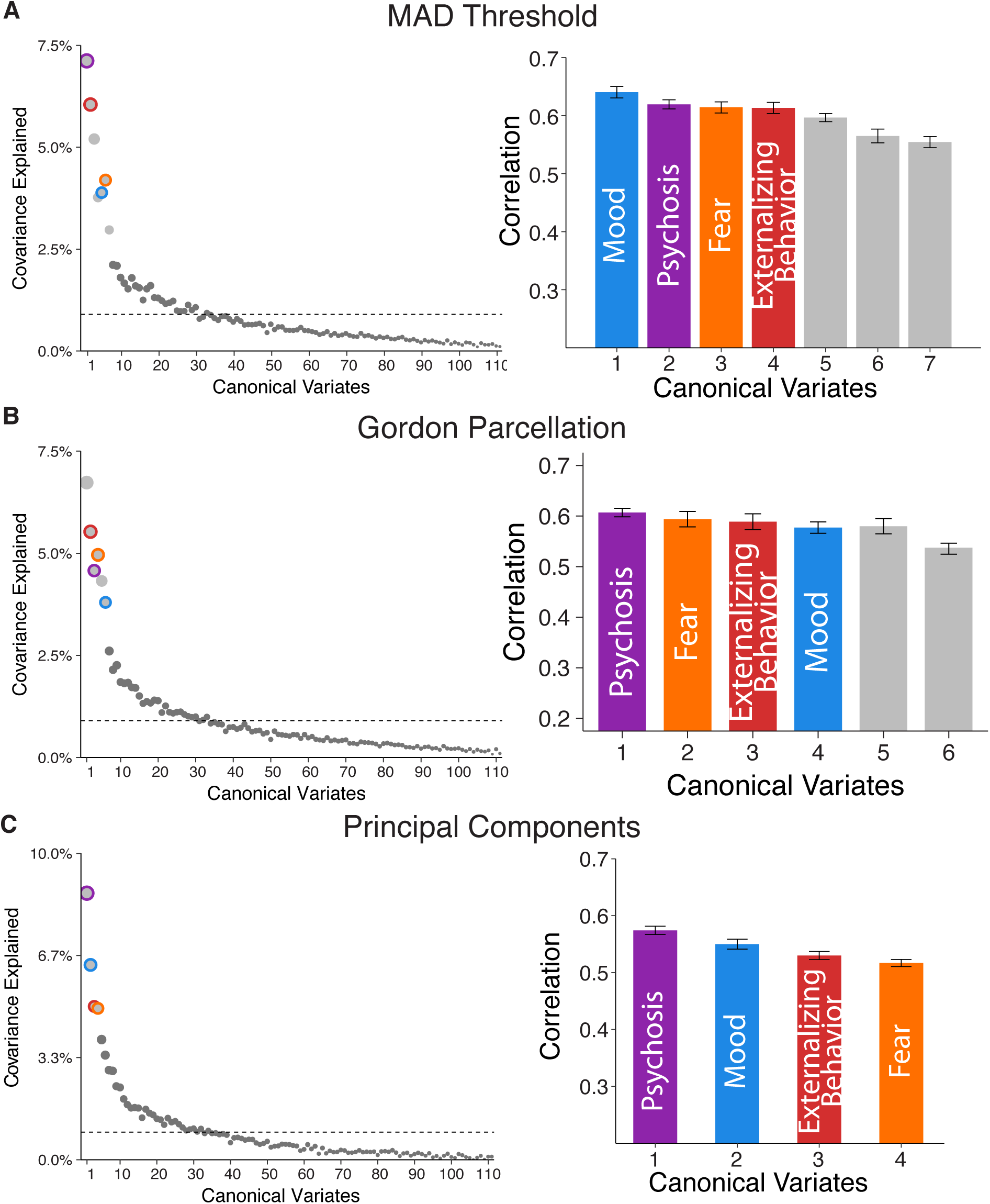
Patterns of canonical variates were robust to methodological choices. We found four canonical variates based on covariance explained and correlation across methodological choices, including **(a)** the number of features entered into the analysis (edges with top 5% variance based on MAD), **(b)** an alternative parcellation (Gordon et al.^99^), and **(c)** using alternative techniques of dimensionality reduction (the first 111 principal components). Dashed line marks the average covariance explained. Corresponding variates on the right panels are circled in the left. Error bars denote standard error.

**Supplementary Figure 6.**
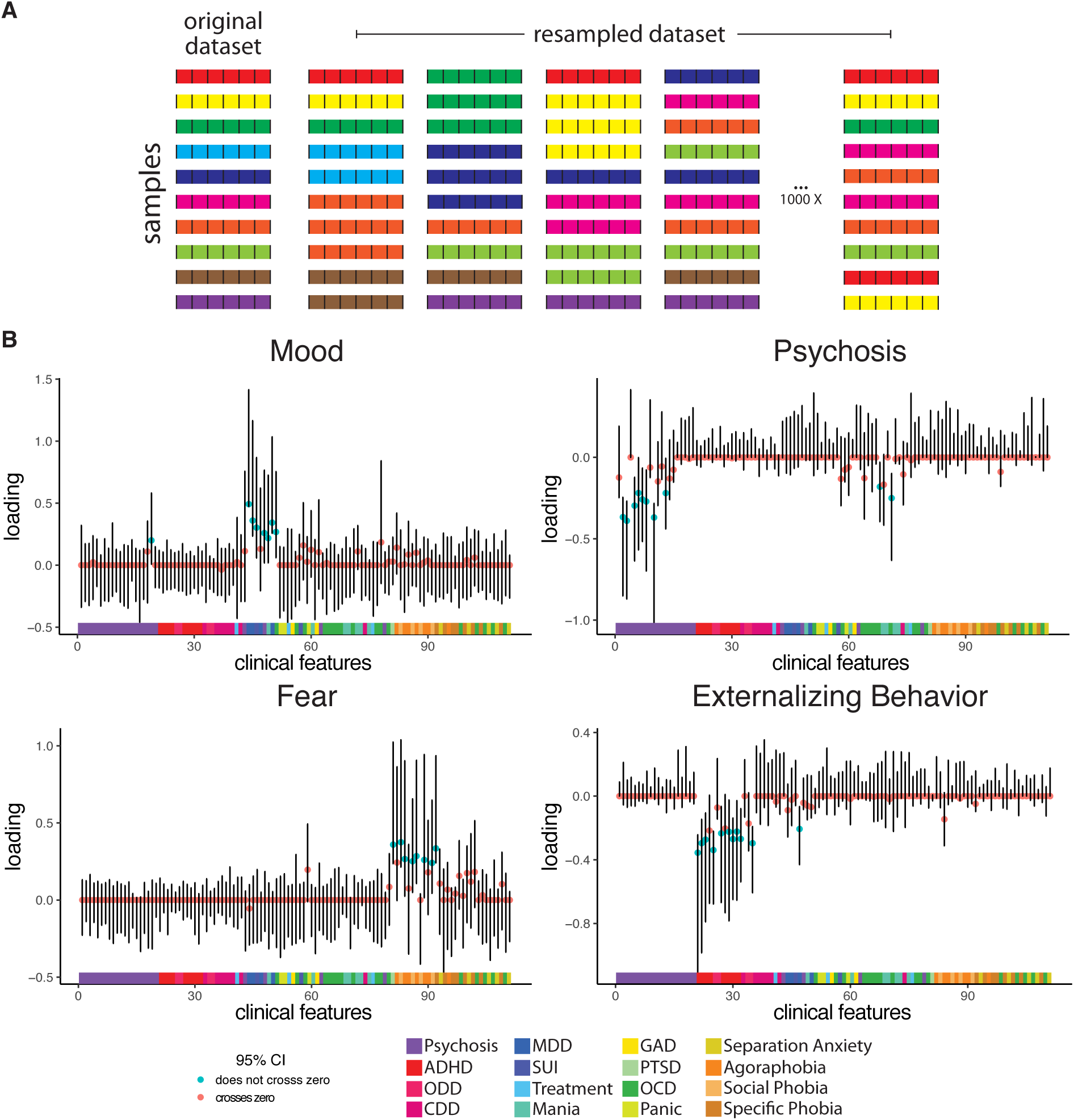
Resampling procedure to identify stable features contributing to each linked dimension. **(a)** Schematic of the resampling procedure. In each sample, two-thirds of the discovery dataset was first randomly selected. The sample size was completed to be the same as the original by replacing with those already selected. **(b)** Resampling distribution for clinical features in each linked dimension. Each bar represents the 95% confidence interval. DSM categories to which each symptom item belongs are shown.

**Supplementary Figure 7.**
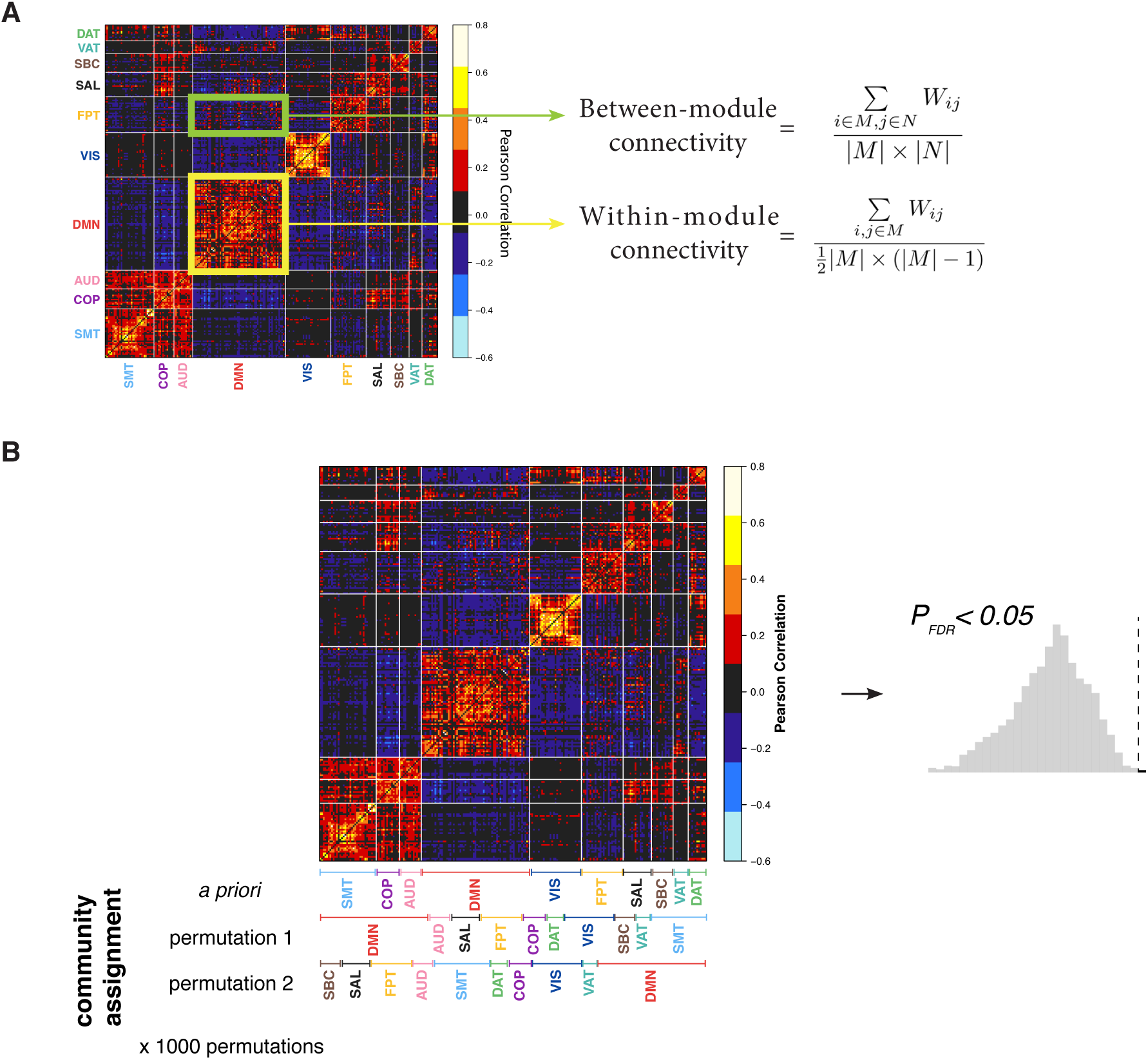
Network module analysis. **(a)** Summarizing loadings on a *between*- and *within*-network basis using *a priori* community assignment from the parcellation of Power et al.^100^ **(b)** Schematic for generating null model for modular analysis. Community membership was randomly assigned to each node while controlling for community size. Mean *between*- and *within*-module loadings were then calculated based on these permuted modules, which we used to assess the statistical significance by comparing the orginal values against the null distribution.

**Supplementary Figure 8.**
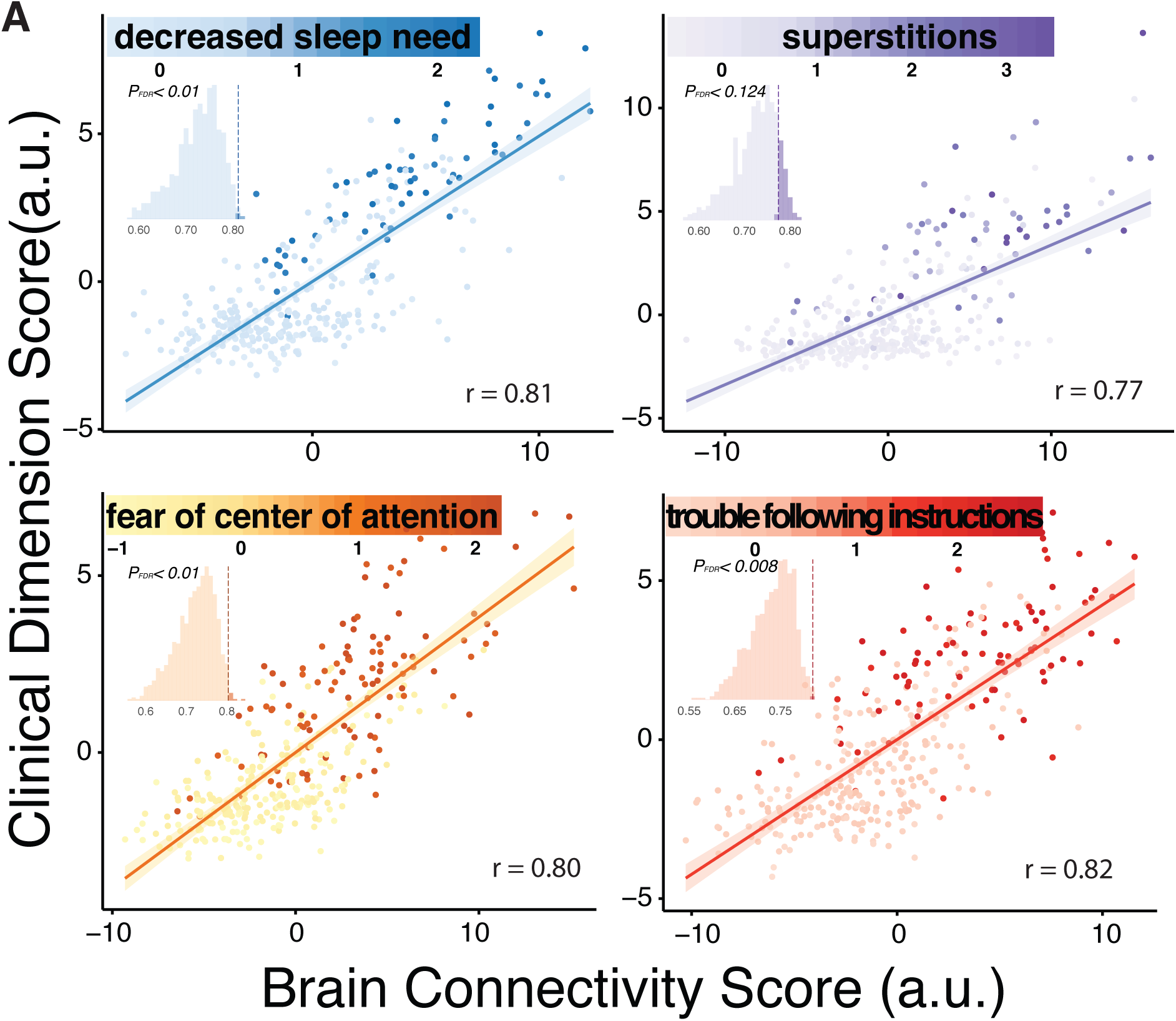
Canonical variates in the replication sample accord with those found in the discovery sample. **(a)** Scatter plots of brain and clinical scores (linear combinations of functional connectivity and psychiatric symptoms, respectively) demonstrate the correlated multivariate patterns of connectomic and clinical features. Colored dots in each panel indicate the severity of a representative clinical symptom that contributed the most to this canonical variate. Each insert displays the null distribution of sCCA correlation by permutation testing. Dashed line marks the actual correlation.

### Supplementary Tables

**Supplementary Table 1.** Demographics in each sample

|  |  | Discovery | Replication | Total |
| --- | --- | --- | --- | --- |
| n |  | 663 | 336 | 999 |
| Sex | Male | 293 | 155 | 448 |
|  | Female | 370 | 181 | 551 |
| Race | White | 306 | 153 | 459 |
|  | Black | 286 | 141 | 427 |
|  | Other | 71 | 42 | 113 |
| Age | 8-10 | 70 | 40 | 110 |
|  | 11-13 | 125 | 63 | 188 |
|  | 14-16 | 195 | 102 | 297 |
|  | 17-19 | 206 | 100 | 306 |
|  | 20-22 | 58 | 30 | 88 |
|  | >22 | 9 | 1 | 10 |
|  | mean | 15.82 ± 3.32 | 15.65 ± 3.32 | 15.76 ± 3.32 |

**Supplementary Table 2.** Correlations of loadings between covariate-regressed and non-regressed features

|  | Connectivity | Symptoms |
| --- | --- | --- |
| Mood | 0.73 | 0.54 |
| Psychosis | 0.95 | 0.88 |
| Fear | 0.70 | 0.35 |
| Externalizing behavior | 0.98 | 0.97 |

Loadings of both connectivity and clinical features across dimensions were highly correlated between input data that had age and sex regressed out of and those that had not. All correlations were statistically significant ( $P_{FDR} < 0.001$ ).

**Supplementary Table 3.**
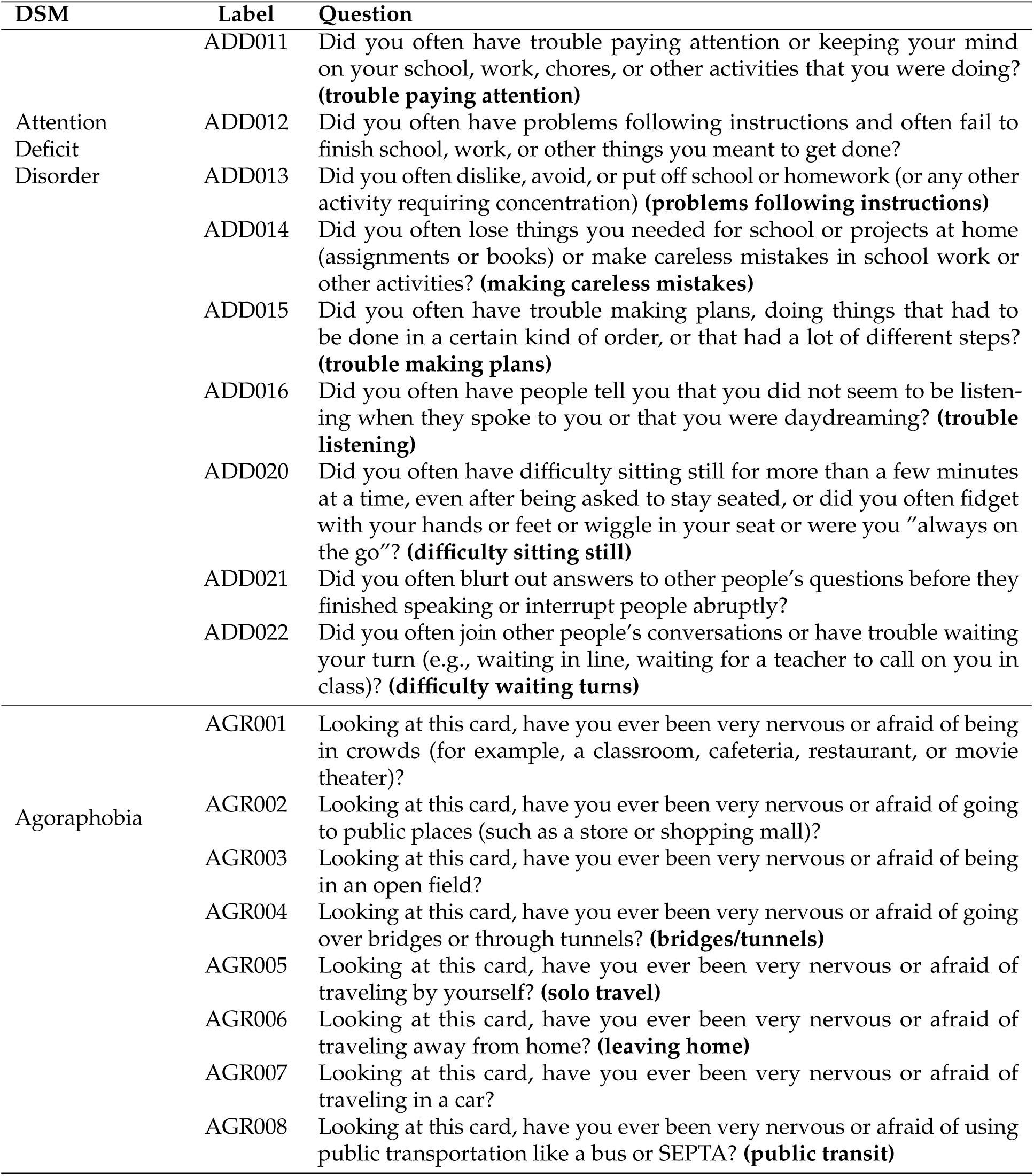
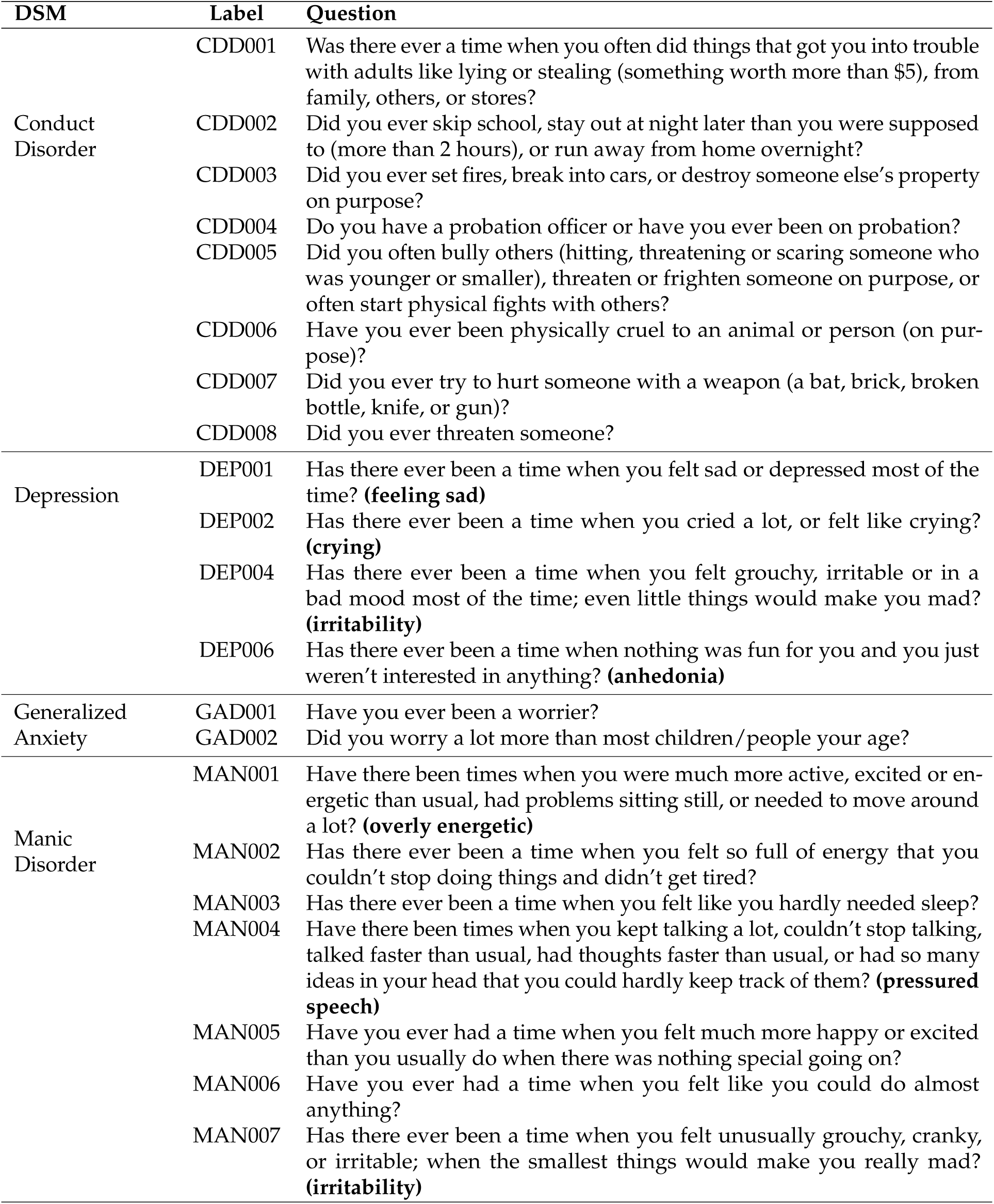
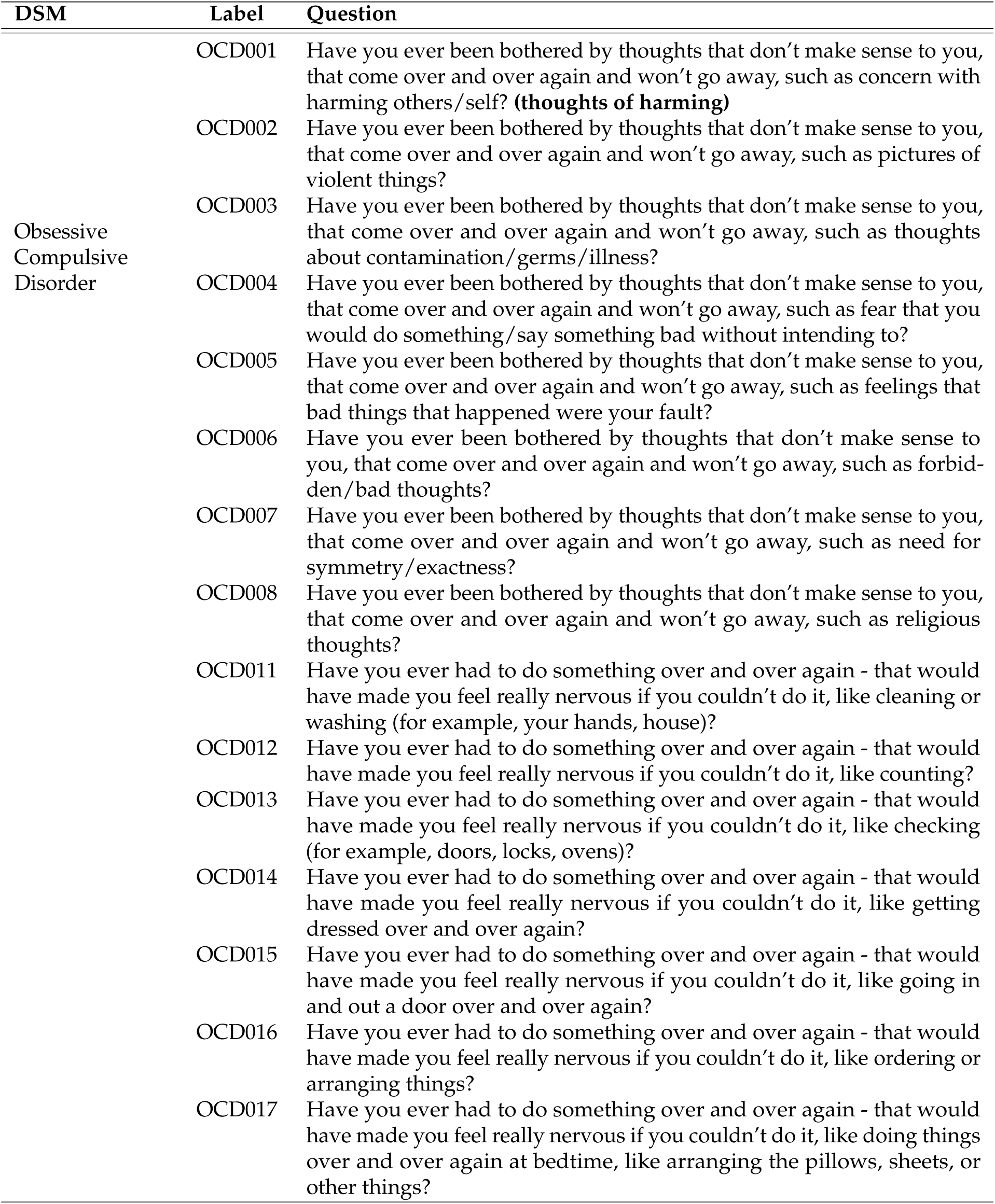
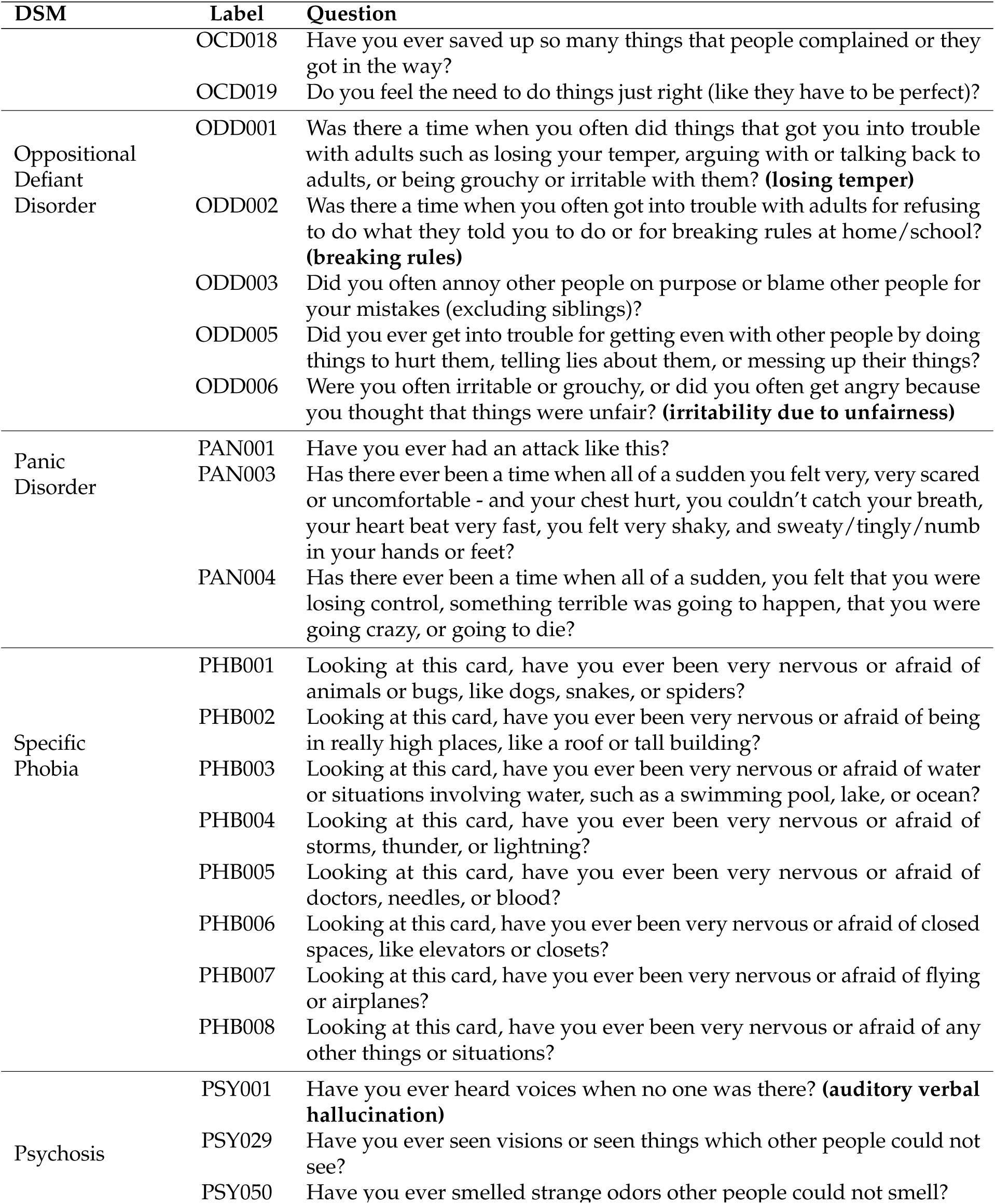
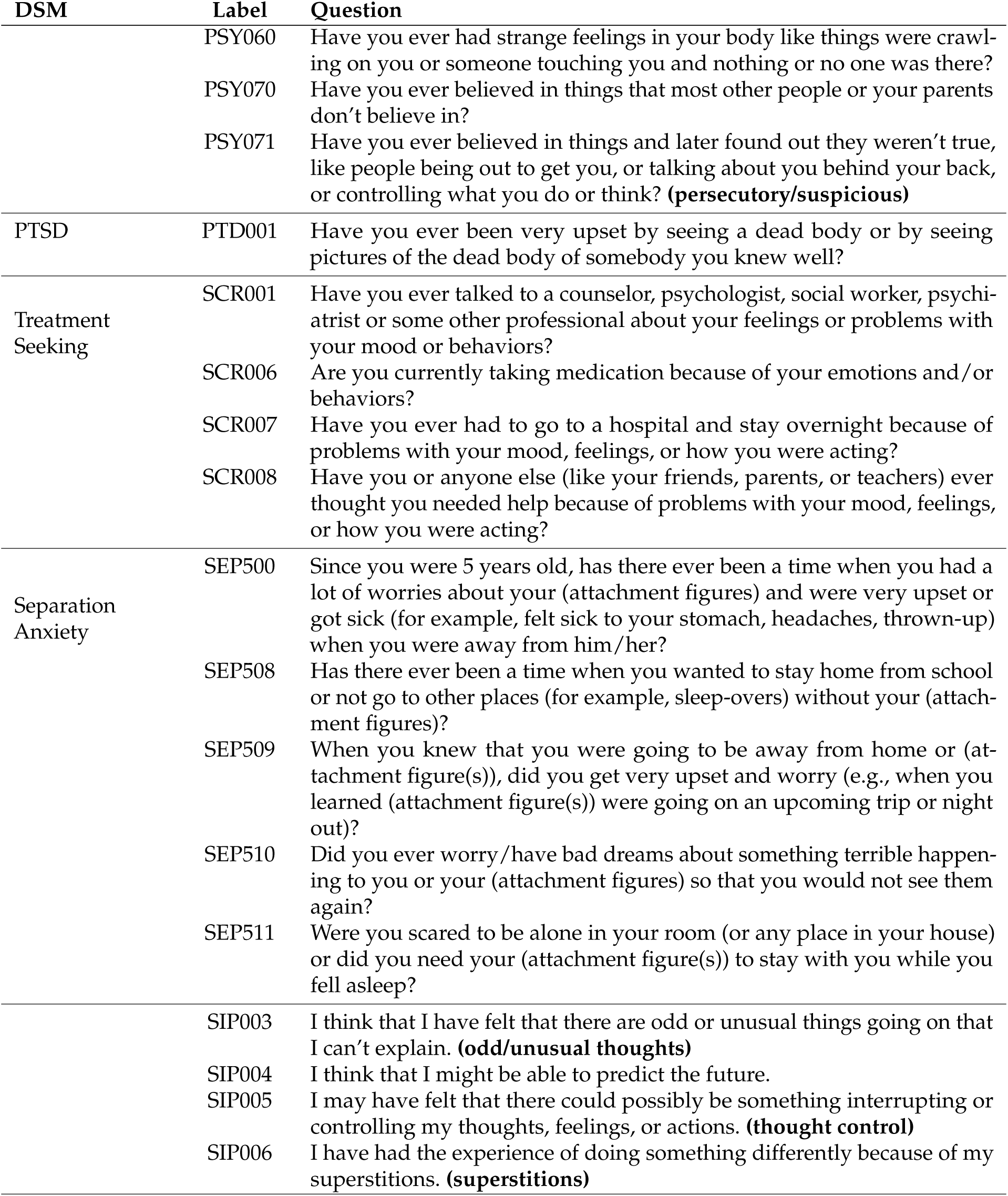
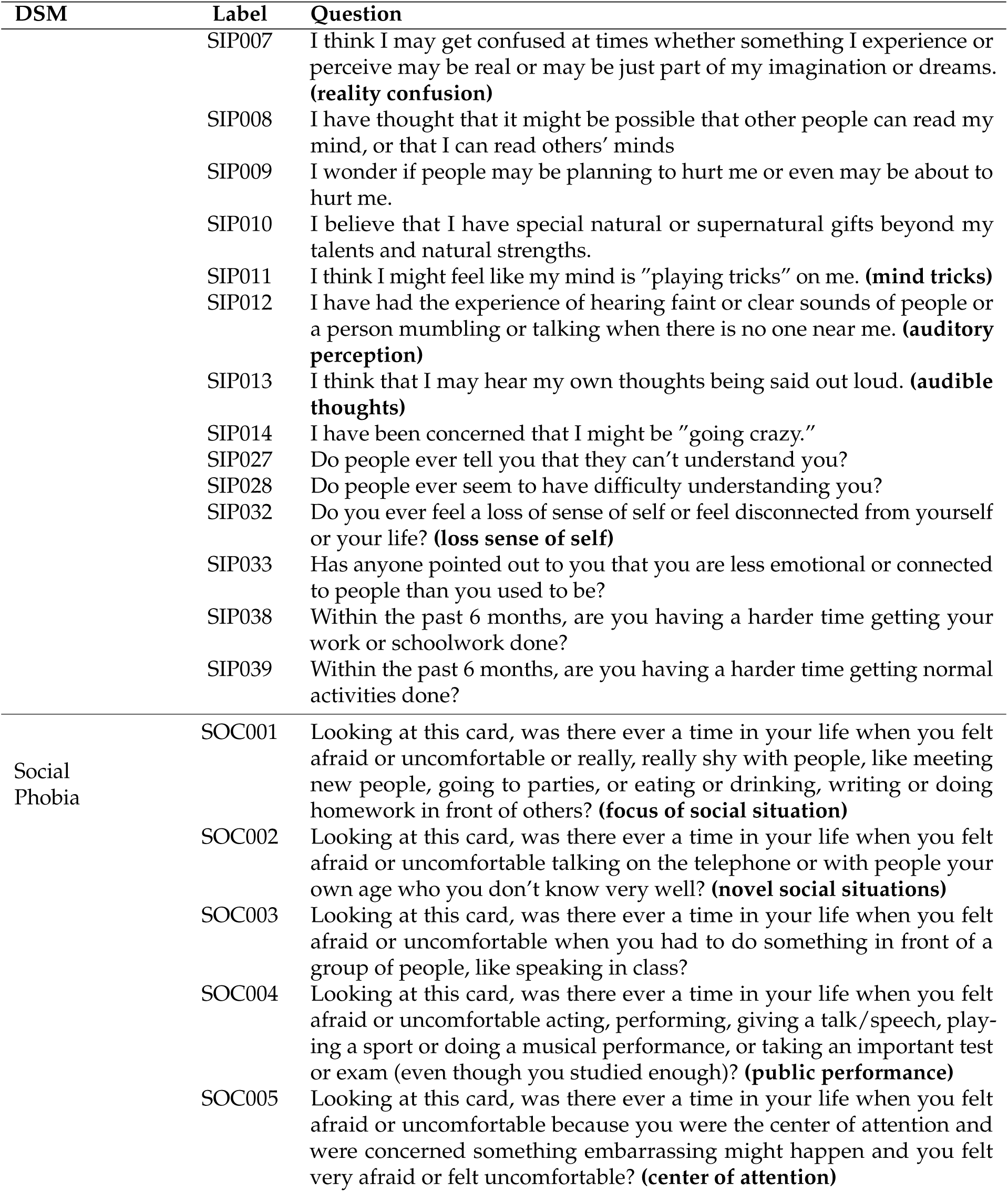
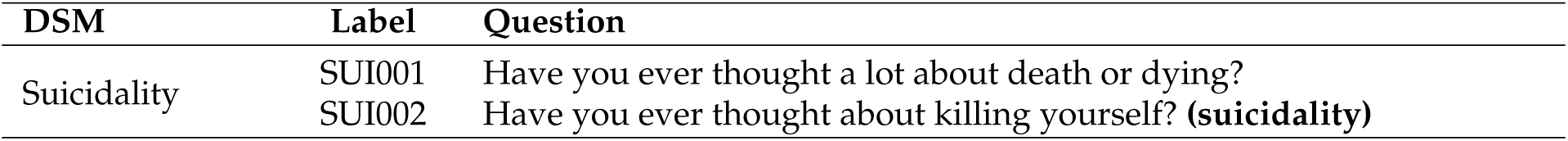
Clinical Assessment

